# SWEET11 and SWEET12 transporters function in tandem to modulate sugar flux in Arabidopsis: An account of the underlying unique structure–function relationship

**DOI:** 10.1101/2022.03.26.485957

**Authors:** Urooj Fatima, D. Balasubramaniam, Wajahat Ali Khan, Manu Kandpal, Jyothilakshmi Vadassery, Arulandu Arockiasamy, Muthappa Senthil-Kumar

## Abstract

Sugar will eventually be exported transporters (SWEETs) have been identified as a unique class of sugar efflux transporters in all biological kingdoms. AtSWEET11 and AtSWEET12 in Arabidopsis act synergistically to perform distinct physiological roles, particularly in apoplasmic phloem loading, seed filling, and sugar level alteration at the site of pathogen infection. Plasma membrane-localized AtSWEET11 and AtSWEET12 transporters exclusively facilitate sucrose transport along the concentration gradient. This article examines the sucrose binding pocket of AtSWEET11 and AtSWEET12 using docking studies, and how they act synergistically in various functions throughout plant development and during abiotic and biotic stresses. Further, we highlight the phylogenetic and the *in-silico* analyses of *AtSWEET11* and *AtSWEET12* orthologs from 39 economically important plant species that could provide new platforms for future studies on sugar allocation mechanisms across the different plant families. In-depth understanding of these transporters and their molecular regulatory mechanisms could be harnessed for crop improvement and crop protection.

## Introduction

Plant sugar will eventually be exported transporter (SWEET) proteins were initially identified in the model organism *Arabidopsis thaliana* (Chen et al., 2010). In total, 17 members were identified in Arabidopsis, classified into four different subclades (Chen et al., 2010; Feng and Frommer, 2015; Breia et al., 2021). All members of Clade III including AtSWEET11 and AtSWEET12 were characterized as sucrose efflux transporters using forster resonance energy transfer (FRET) based sucrose sensors expressed in human embryonic kidney (HEK293T) cells, yeasts, and time-dependent efflux of [^14^C] sucrose in *Xenopus* oocytes (Chen et al., 2012). AtSWEETs are bidirectional transporters that facilitate the diffusion of sucrose molecules down the concentration gradient and adapt a uniporter transport mechanism as their transport activity is pH independent. Kinetic studies of AtSWEET12 (Km for sucrose uptake and efflux was ∼70mM and 10 mM respectively) revealed that SWEETs are low affinity sucrose transporters (Chen et al., 2012). AtSWEET11 and AtSWEET12 share almost 88% amino acid similarity (Chen et al., 2012) and both were shown to exhibit the substrate flexibility as they transported glucose and fructose in addition to sucrose (Le Hir et al., 2015). These SWEET proteins are heptahelical transmembrane (TM) transporters with two internal parallel triple-helix bundles (THB) that are interconnected by the nonconserved helix TM4 (Han et al., 2017; Tao et al., 2015). The two THB domains have a twofold rotation symmetry perpendicular to the membrane plane and have a characteristic 1-3-2 and 5-7-6 topological arrangement (Han et al., 2017; Anjali et al., 2020).

Among the members of *AtSWEETs, AtSWEET11* and *AtSWEET12* are particularly known to be expressed in all tissues, including the leaves, roots, seeds, siliques, and flowers (Chen et al., 2012). In Arabidopsis, AtSWEET11 and AtSWEET12 participate in pivotal processes such as phloem loading (Chen et al., 2012), xylem development (Le Hir et al., 2015), and seed filling (Chen et al., 2015). This demonstrates that AtSWEET11 and AtSWEET12 are crucial transporters required in major developmental and physiological processes. Besides, these two transporters optimize the sugar flux in response to varying environmental conditions. AtSWEET11 and AtSWEET12 have been shown to have role during interactions with pathogens (Chen et al., 2010; Gebauer et al., 2016; Walerowski et al., 2018). Nevertheless, the dynamic role of these two transporters in multiple integrated aspects of plant physiology has raised considerable interest in their structure and molecular regulatory mechanisms. In this article, we probe the structural aspects of sucrose binding pockets from AtSWEET11 and AtSWEET12 with the help of molecular docking and molecular dynamics simulations. We also draw critical insights about the physiological functions of AtSWEET11 and AtSWEET12 and provide in-depth analyses of their orthologs from economically important crops, as well as the possible mechanisms of regulation.

## Results and Discussion

### AtSWEET11 and AtSWEET12 structures and their substrate-binding pockets

The crystal structures of OsSWEET2b (from *Oryza sativa*) (Tao et al., 2015) and AtSWEET13 (Han et al., 2017) have recently been determined. The TM regions of AtSWEET11 and AtSWEET12 were homology modelled using the Robetta server (Yang et al., 2020) (**Figure 1 and Supplementary Figure 1**). The TM regions of AtSWEET11 and AtSWEET12 share similar structural features with OsSWEET2b and AtSWEET13, as indicated by low root mean square deviation (RMSD) upon superposition (**Supplementary Figure 2**). Both AtSWEET11 and AtSWEET12 models consist of seven TM helices (TMs 1–7), with N-terminal THB (TMs 1–3) and C-terminal THB (TMs 5–7) interconnected by TM4 (**Figure 1A, B, D and E**). The homology models of AtSWEET11 and AtSWEET12 are in cytoplasmic open conformation (**Supplementary Figure 2 and Figure 1G**). The TM region for AtSWEET11 and AtSWEET12 from the AlphaFold Protein Structure Database (Jumper et al., 2021 and Varadi et al., 2021) were superposed (Cα atoms) with respective models reported in this article and found to be in cytoplasmic open conformation with no significant RMSD. The cytoplasmic C-terminal domain (CTD) of AtSWEET11 (220–289 amino acids) and AtSWEET12 (220–285 amino acid) could not be modeled with accuracy due to lack of homologous structures.

**Figure 1.**
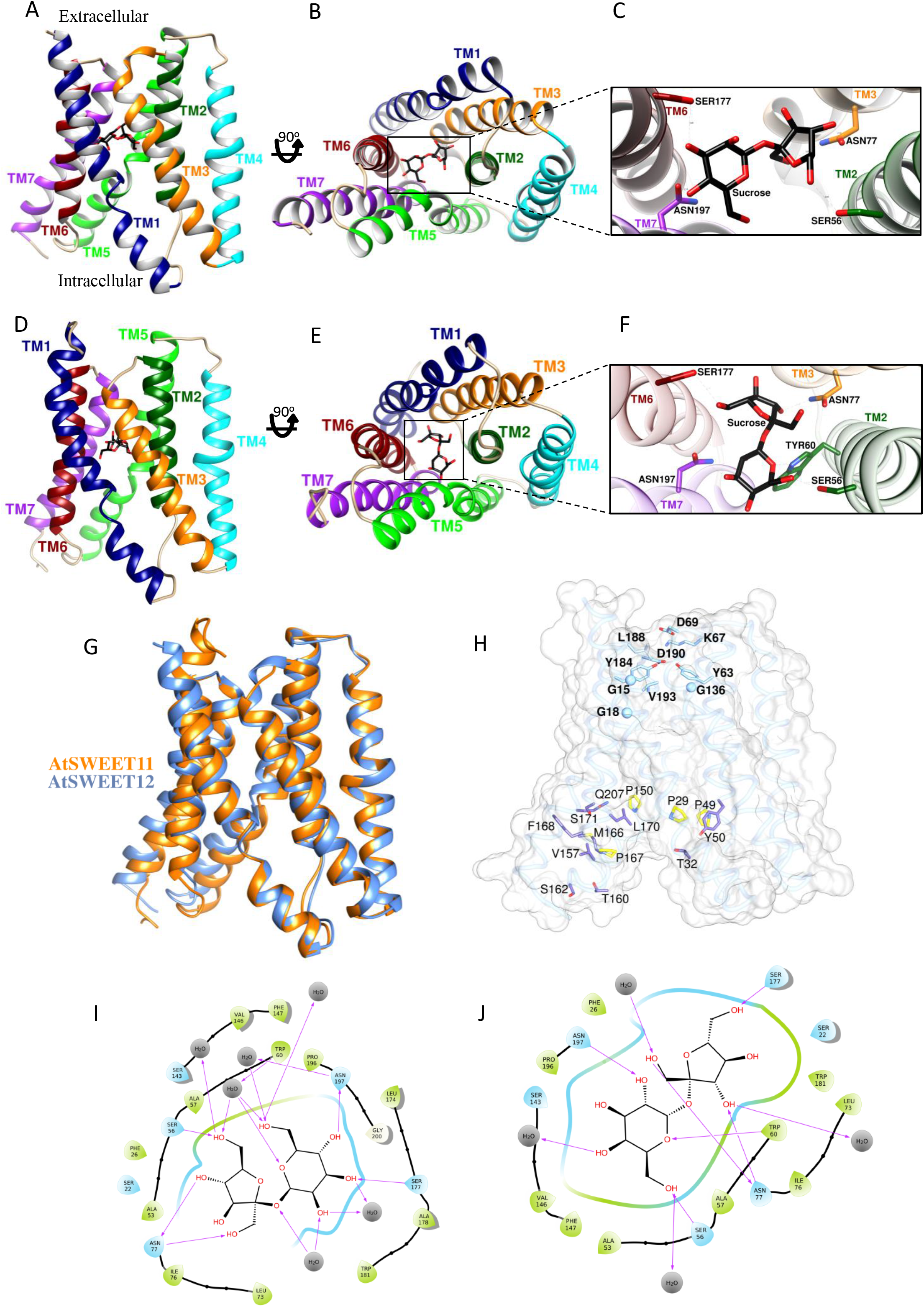
Protein structures of AtSWEET11 and AtSWEET12 and key residues involved in sucrose transport. **A–J**, In silico homology models of AtSWEET11 and AtSWEET12. The stable docked poses of the SWEET–sucrose complex after molecular dynamics (MD) simulations are shown. The transmembrane (TM) helices are colored (TM1–TM7), and sucrose is represented as a black stick. Ribbon representation of AtSWEET11 in parallel view from the membrane **(A)**, in cytoplasmic view **(B)**, in a close-up view of sucrose in the central cavity with interacting residues **(C)**. Ribbon representation of AtSWEET12 in parallel view from the membrane **(D)**, in cytoplasmic view **(E)**, in a close-up view of sucrose in the central cavity with interacting residues **(F). (G)**, Structural superposition of AtSWEET11 (cornflower blue) and AtSWEET12 (orange). The root mean square deviation (RMSD) between the structures is 1.48 Å. **(H)**, The residues of the periplasmic gate and cytosolic gate in AtSWEET11 (are represented as a transparent gray surface and blue ribbon), respectively. The residues of the periplasmic gate are shown in cyan. The proline tetrad ring from the cytosolic gate is shown in yellow. The other putative residues that might be involved in cytoplasmic gating during the transport cycle are shown in purple: T32, Y50, V157, T160, S162, M166, F168, L170, S171, and Q207. The sucrose-interacting diagram for the complex in **(I)** AtSWEET11 and **(J)** AtSWEET12. For more details, see Supplementary Figure 1 and 3.

Molecular docking followed by molecular dynamics (MD) simulations for 500 ns and 1000 ns, respectively were performed in order to probe the sucrose binding site using site map (Schrödinger’s suite) (**Supplementary Figure 1 and 3**). For both AtSWEET11 and AtSWEET12, the best scoring sites predicted at the central cavity and in the middle of the TM region (**Supplementary Figure 3**) were used to dock the sucrose (**Figure 1C and F**), which is unbiased by the already known crystal structure of AtSWEET13-dCMP complex. This allows alternate access to the sucrose molecule from either side of the membrane during transport. Interestingly, in both AtSWEET11 and AtSWEET12, the residues S22, S56, W60, N77, N197, W181, S177, V146, and S143 interact with sucrose, with equivalent residues interacting with dCMP in the reported AtSWEET13 structure (**Figure 1I and J, Supplementary Table 1**). The additional sucrose interacting residues in AtSWEET11 and AtSWEET12 from the molecular simulation is represented as 2D-Ligand interaction diagram (Figure 1I and Figure 1J). Among the residues present in the sucrose binding site, Asn77 and Asn197 are highly conserved across all AtSWEETs (except AtSWEET6, wherein Asn77 is replaced by serine) and OsSWEET2b. Multiple sequence alignment of AtSWEET11, AtSWEET12, and AtSWEET13 with OsSWEET2b reveals 61 conserved residues, whereas only 22 are conserved across all the AtSWEETs 1-17. Notably, T159 in AtSWEETs is replaced by serine in OsSWEET2b (**Supplementary Figure 4**). In both AtSWEET11 and AtSWEET12, Y63, K67, D69, and D190 form the putative extracellular gate (**Figure 1H**). The Tyr–Asp pair (Tyr63 and Asp190) that forms the periplasmic gate is highly conserved among SWEETs. However, the interaction of this pair with a Gln132 (OsSWEET2b) or Glu131 (AtSWEET13) is lost due to replacement with Ala in AtSWEET11 and AtSWEET12. Other residues, namely, Y184, V193, and L188, may also form a luminal gate, as mutations in corresponding residues have abolished the transport activity in AtSWEET13 and OsSWEET2b. Of the remaining conserved residues that can form the putative intracellular gate, the proline tetrad (P29 from TM1, P49 from TM2, P150 from TM5, and P167 from TM6) plays a major role in alternating access mechanism of sucrose transport by inducing concerted structural arrangements in the TM helices (**Figure 1H**). Replacing any one of these prolines with alanine in AtSWEET1 abolishes the transport activity (Tao et al., 2015). When adjacent residues to these conserved prolines were mutated to alanine in AtSWEET1, it resulted in reduced glucose transport (Tao et al., 2015), indicating the importance of these residues in the transport cycle. The other conserved residues that form the putative intracellular gate are shown in Figure 1H. When co-expressed with a wild type transporter, mutation of V188A in AtSWEET1 in the extrafacial gate showed complete transport inhibition while P23T in the cytosolic gate played an allosteric role, both displaying a dominant-negative effect in substrate transport (Han et al., 2017; Tao et al., 2015; Xuan et al., 2013).

The oligomeric state of SWEETs, including AtSWEET11 and AtSWEET12, remains obscure. The crystal structure of OsSWEET2b is trimeric (Tao et al., 2015). While the crystal structure of AtSWEET13 is monomeric, multi-angle light scattering with size-exclusion chromatography (SEC-MALS) and single-molecule fluorescence resonance energy transfer (FRET) studies have suggested it to be a dimer (Han et al., 2017). The finding of the cytoplasmic CTD of AtSWEET13 alone forming a dimer (Han et al., 2017) needs further experimental evidence and the involvement of the CTD in oligomerization and the regulation of substrate transport warrants further exploration.Taken together, these studies suggests that the oligomerization of SWEETs is necessary for functional activity which requires further evidence from structure-function studies of native SWEETs from different plant species.

### Role of AtSWEET11 and AtSWEET12 transporters: redundant or exclusive?

During apoplasmic phloem loading, sucrose produced in mesophyll cells is transported into the mesophyll apoplasm, from where it moves towards the phloem (Giaquinta, 1983; Ayre, 2011). AtSWEET11 and AtSWEET12 proteins are localized on the plasma membrane of the phloem parenchyma cells in leaf tissue, where they are involved in the translocation of sucrose from mesophyll cells to the phloem apoplasm (Chen et al., 2012). The sucrose effluxed into the phloem apoplasm is transported to the sieve element–companion cell (SE/CC) complex with the help of the sucrose transporter AtSUC2 against a concentration gradient (Srivastava et al., 2008; 2009a,b). Chen et al. (2012) showed that plants carrying mutations in both *AtSWEET11* and *AtSWEET12* showed moderate growth defects and accumulated excessive sugar in the leaves due to blockage in the sugar translocation pathway; this suggests their critical role in the phloem-loading process (**Figure 2A**). However, single mutants of either of these genes did not affect the plant (Chen et al., 2012), implying a redundant function of AtSWEET11 and AtSWEET12 in phloem loading.

**Figure 2.**
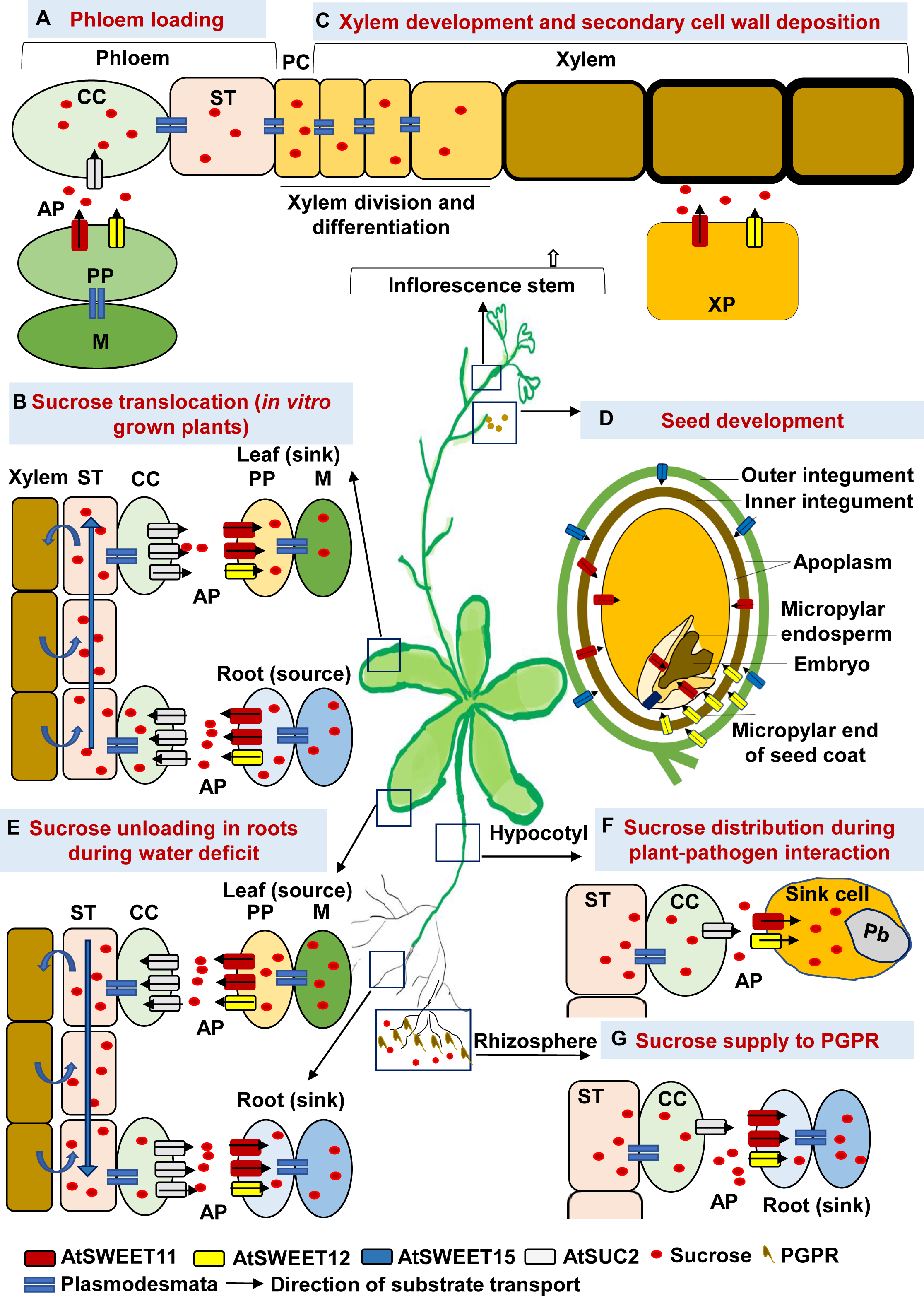
AtSWEET11 and AtSWEET12 localization, involvement in the sucrose transport pathway, and related physiological functions and stress responses in Arabidopsis. AtSWEET11 and AtSWEET12 transporters are reported to play a major role in the roots, hypocotyl, leaf/stem (tracheid), and seeds. **A**, During apoplasmic phloem loading, phloem parenchyma-localized AtSWEET11 and AtSWEET12 transport sucrose from the mesophyll cells to the phloem apoplast. Next, AtSUC2 transports the sucrose from the apoplast to the sieve element–companion cell complex (Chen et al., 2012). **B**, In *in vitro* grown plants with exogenous carbon supply, the roots act as a source tissue, and the leaves become the sink. The directionality of sucrose transport by AtSWEET11 and AtSWEET12 gets reversed according to the sucrose concentration gradient, i.e., from the root (source) to the leaf tissues (sink) (Papaioannou et al., 2018). **C**, During xylem development, AtSWEET11 and AtSWEET12 are localized in xylem parenchyma cells in the inflorescence stem and facilitate secondary cell wall formation by delivering carbon sources to the developing xylem cells in the inflorescence (Le Hir et al., 2015). **D**, During seed filling, AtSWEET15 localized in the outer integument transports sucrose into the apoplast. AtSWEET11 localized in the inner integument facilitates sucrose supply to the developing embryo from the micropylar endosperm. AtSWEET12 localized in the micropylar end of the seed coat facilitates sucrose transport to the seed coat region. Together, these three transporters, among others, are involved in sink-drawing ability during seed development/grain filling (Chen et al., 2015). **E**, During water deficit conditions, AtSWEET11, AtSWEET12, and AtSUC2 are involved in drawing more sucrose to the root cells. Under water stress, these transporters accumulate in the roots and unload sucrose from the apoplast to the sink cells in the roots and facilitate root growth by allocating more sucrose from the leaves to the roots (Durand et al., 2016). **F**, AtSWEET11 and AtSWEET12 facilitate sugar delivery towards the pathogen (here, *Plasmodiophora brassicae* at the site of infection in the hypocotyl region) (Walerowski et al., 2018). **G**, AtSWEET11 and AtSWEET12 participate in phloem unloading of sucrose, especially to the lateral roots. These transporters control the sugar supply from the shoot to the root and then distribute sugars to plant growth-promoting rhizobacteria (PGPR, i.e., here, *Pseudomonas simiae* WCS417r in the rhizosphere region) (Desrut et al., 2020). CC: companion cell; ST: sieve tubes; PP: phloem parenchyma; M: mesophyll cells; PC: pro-cambium cells; XP: xylem parenchyma; AP: apoplast; Pb; *Plasmodiophora brassicae*

AtSWEET11 and AtSWEET12 are also involved in long-distance translocation of sucrose in both the source and sink of plants grown *in vitro* (**Figure 2B**) (Papaioannou et al., 2018). *In vitro* grown plants with exogenous carbon supply, where the roots act as source tissue, showed *AtSWEET11* and *AtSWEET12* expression within both leaf and root tissues; however, more pronounced expression was observed in the roots (Papaioannou et al., 2018). The study suggests that the directionality of sucrose transport by AtSWEET11 and AtSWEET12 can be reversed in accordance to the sucrose concentration gradient. Besides, the study showed that the transgenic with the single mutant *atsweet12* and *AtSWEET11* overexpression translocated more sucrose from the source (root tissues) to the sink (leaf tissues). However, this was not observed in the reverse case, i.e., *atsweet11* mutant with *AtSWEET12* overexpression (Papaioannou et al., 2018). In other words, in the absence of AtSWEET12, sucrose transport is overtaken by AtSWEET11 but not vice versa; hence, AtSWEET11 is crucial for sucrose transport. Surprisingly, in the same study, *ex vitro* grown plants, in which the leaves acted as the source tissue, showed higher expression of *AtSWEET11* than that of *AtSWEET12*. Similarly, in another study, endogenously supplied sucrose—by providing high light conditions—showed contrasting effects on the expression of *AtSWEET11* and *AtSWEET12* (Wei et al., 2020). Thus, in the presence of high endogenous sucrose levels in leaf tissues, *AtSWEET11* expression is upregulated, but *AtSWEET12* expression is not. Taken together, these studies argue against the redundant roles of AtSWEET11 and AtSWEET12 under certain conditions and hint at the dominant role of AtSWEET11 in sucrose transport function.

AtSWEET11 and AtSWEET12 are expressed in the phloem cells of the flower stem and might contribute to the transport of sugars from the phloem to provide nutrition to the adjacent stem tissues (Chen et al., 2012; Le Hir et al., 2015). Besides their phloem-loading functions, AtSWEET11 and AtSWEET12 facilitate secondary cell wall formation by delivering carbon sources to developing xylem cells (Le Hir et al., 2015) (**Figure 2C**). Phenotypic and anatomical characteristics revealed reduced stem diameter and xylem and phloem areas in single *atsweet11* and double *atsweet11;12* mutants, but no distinguishable features were observed in *atsweet12* mutant. These observations established that the mutation of *AtSWEET11* has a dominant effect on the phenotype compared to the mutation of *AtSWEET12* alone. However, the double mutant phenotype is more prominent than that of single mutants (Le Hir et al., 2015), suggesting that AtSWEET11 and AtSWEET12 transporters work in tandem to regulate sugar transport.

During seed filling, sucrose is transported through the phloem to the maternal seed coat and from the seed coat to provide nutrition to the developing embryo (Stadler et al., 2005). AtSWEET11, AtSWEET12, and AtSWEET15 are crucial for seed filling in Arabidopsis (Chen et al., 2015) (**Figure 2D**). The *atsweet11;12* double and *atsweet11;12;15* triple mutants showed seed defects, but the single mutants did not. However, the defects in seed phenotype were more severe in *atsweet11;12;15* triple mutants, where the embryo development, seed weight, and starch and lipid content were drastically compromised (Chen et al., 2015). The spatiotemporal localization of these genes at multiple sites in the seed tissue revealed the feeding pathway for the developing embryo. AtSWEET15, which is present in the outer integument, transports sucrose into the apoplast. The localization of AtSWEET11 in the inner integument facilitates sucrose efflux from the inner integument. Together, AtSWEET11 and AtSWEET15 supply sucrose to the developing embryo from the micropylar endosperm. AtSWEET12 accumulation in the micropylar end of the seed coat facilitates sucrose transport to the seed coat region (Chen et al., 2015). Irrespective of their spatiotemporal localization, these transporters translocate sucrose along the concentration gradient driven by the thermodynamically favored transport. While the mutant study indicated the redundant function of these three genes, the spatiotemporal accumulation of AtSWEET11, AtSWEET12, and AtSWEET15 proteins points to the distinct and exclusive role of each of these transporters, working in tandem during the seed development process.

### AtSWEET11 and AtSWEET12 regulate sugar flux in response to fluctuating environments

#### In response to abiotic stimulus

Water deficit conditions alter sugar allocation in the source and sink tissues and also impact the expression of the sugar transporters involved in long-distance sucrose transport. The transcript levels of the *AtSWEET11, AtSWEET12*, and *AtSUC2* genes are highly induced in the leaves of plants under water deficit conditions and these sucrose transporters were predicted to be involved in exporting sucrose accumulated in the leaves to the roots (Durand et al., 2016). In the recent detailed study the phosphorylation of sucrose transporters was shown to direct enhanced root:shoot ratio in plants under drought stress (Chen et al., 2022; Gong and Yang, 2022). In addition, *AtSWEET11, AtSWEET12*, and *AtSUC2* gene expression was also induced in water-stressed roots, indicating the potential role of these transporters during sucrose-unloading processes (Durand et al., 2016; Gong and Yang, 2022) (**Figure 2E**). It is evident that under water deficit conditions, plants sustain their growth and development by allocating more sugars from the leaves to the roots by utilizing these transporters.

Under cold stress, soluble sugars, including sucrose molecules, act as osmoprotectants to maintain cellular integrity (Ruan et al., 2010). Le Hir et al. (2015) showed that *AtSWEET11* and *AtSWEET12* expression is downregulated after cold stress treatment. Cold treatment induces a high level of glucose and fructose accumulation in *atsweet11* and *atsweet11;12* mutants compared to the wild type, and no alteration was seen in the sugar content of *atsweet12* mutants. Moreover, only the double mutant exhibited freezing tolerance (Le Hir et al., 2015), suggesting the synergistic role of both the transporters in regulating the sugar flux inside plant cells during cold stress.

#### In response to biotic stimulus

SWEETs have been shown to be targeted by bacterial effectors to provide nutrients to bacterial pathogens and establish infection in crop plants (Antony et al., 2010; Chen et al., 2010; Chong et al., 2014; Cox et al., 2017; Li et al., 2012). Chen et al. (2010) reported the induced transcript levels of *AtSWEET11* and *AtSWEET12* genes during infection with biotrophic, hemibiotrophic, and necrotrophic pathogens, suggesting the modulation of these transporters to provide nutrients to pathogens. However, the findings of Gebauer et al. (2016) did not support the role of AtSWEET11 and AtSWEET12 transporters in pathogen nutrition in Arabidopsis after infection with the fungal hemibiotroph *Colletotrichum higginsianum*. Their study showed that *atsweet11;12* double mutants, but not the single mutants, were resistant to *C. higginsianum* due to sugar-mediated priming of the salicylic acid pathway. However, the strong accumulation of AtSWEET12-YFP fusion proteins at the localized infection sites, but not of AtSWEET11-YFP fusion proteins, does not support the redundant roles of AtSWEET11 and AtSWEET12 transporters. The Arabidopsis interaction with the protist biotroph *Plasmodiophora brassicae* established the role of AtSWEET11 and AtSWEET12 transporters in facilitating sugar delivery towards pathogens at the infection site (Walerowski et al., 2018). AtSWEET11 and AtSWEET12 expression levels were highly induced in the phloem cells of the infected hypocotyl tissues. The *atsweet11;12* double mutants, but not the single mutants, were defective in supplying sugars to the pathogen, thereby impairing gall development and retarding disease progression (Walerowski et al., 2018) **(Figure 2F)**. However, Li et al. (2018) showed that the progression of disease development by *P. brassicae* was delayed in the *atsweet11* single mutants, which again contradicts the functional redundancy of AtSWEET11 and AtSWEET12 transporters during *P. brassicae* infection.

Transcript expression of *AtSWEET11* and *AtSWEET12* was highly induced after inoculation of Arabidopsis with plant growth-promoting rhizobacteria (PGPR) strain *Pseudomonas simiae* WCS417r, and the beneficial effect of *P. simiae* was lost in Arabidopsis *atsweet11;12* double mutants (Desrut et al., 2020). This suggests that AtSWEET11 and AtSWEET12 could possibly function in controlling the sugar supply from the shoot to the root and its distribution to the PGPR, which might positively impact the plant–PGPR interaction (**Figure 2G**). Moreover, histochemical analysis using the GUS reporter assay also confirmed the function of AtSWEET12 in phloem unloading of sucrose to the primary and the lateral roots (Desrut et al., 2020). Also, AtSWEET11 and AtSWEET12 were shown to play a role in blocking the sugar supply to the propagating hyphae of *Trichoderma harzianum*. The host restricts sugar availability in the root apoplast by downregulating *AtSWEET11* and *AtSWEET12* genes for phloem unloading and by upregulating *AtSUC2* and *AtSWEET2* genes for sugar uptake inside the root cells (Rouina et al., 2021). Similarly, AtSWEET12 was shown to suppress multiplication of different species of *Pseudomonas syringae* by restricting sucrose availability to these foliar bacterial pathogens in the leaf apoplast. The study traced the AtSWEET11-mediated sucrose flux to be modulated through AtSWEET12 via plasma membrane targeting and an oligomerization-dependent regulatory mechanism in Arabidopsis. This also indicates the exclusive role of AtSWEET12 in suppressing bacterial multiplication and the role of AtSWEET11 in supplying sugars to bacterial pathogens in the apoplast (Fatima and Senthil-Kumar, 2021).

Nevertheless, focused experiments are required to further validate the exclusive or redundant role of these transporters in phloem unloading and during plant–pathogen interactions.

### Analysis of *AtSWEET11* and *AtSWEET12* orthologs from different plant species

The potential orthologs of *AtSWEET11* and *AtSWEET12* were identified by comparing the protein sequence against the proteomes of 39 economically important plant species from 20 different families (**Supplementary File 1a**). *AtSWEET11* orthologs were found in 24 of the 39 plant species (**Supplementary File 1b**), while *AtSWEET12* orthologs were found in 33 plant species (**Supplementary File 1c**). Interestingly, orthologs exclusive for *AtSWEET11* were found in six different plant species while 12 plant species had orthologs exclusive for *AtSWEET12* (**Supplementary File 1b, c**). The absence of either *AtSWEET11* or *AtSWEET12* orthologs in these plant species implies that during the course of evolution, only one transporter might have been responsible for the sugar transport function. Genome sequence analysis of potential orthologous candidates from different plant species indicates the evolutionary and functional significance of these two transporters across the plant kingdom. The time tree analysis of all 39 species demonstrates a distinct bifurcation between monocots and dicots (**Supplementary Figure 5**). Similarly, the AtSWEET11 and AtSWEET12 orthologs exhibit conservation within dicot and monocot group, except for *Secale cereal* (**Supplementary Figure 6**).

Chromosomal location and gene structure analysis were performed for the identified orthologs of *AtSWEET11* and *AtSWEET12* (**Supplementary File 1b, c**). The gene length for *AtSWEET11* orthologs from different plant species varies from 1,300 to 3,500 bp, with approximately 5–6 exons and 4–6 introns, except in *Solanum lycopersicum*, which has a gene length of approximately 14,000 bp with 12 exons and 12 introns. The orthologs of *AtSWEET12* from different plant species have gene lengths of 1,300–4,000 bp with 6–7 exons and 5–6 introns (**Supplementary Figure 7**). The promoter sequences of *AtSWEET11* and *AtSWEET12* orthologs were analyzed for *cis*-acting regulatory elements. *Cis*-elements involved in plant development, including light response, were present in the promoter region of the *AtSWEET11* and *AtSWEET12* orthologs (**Figure 3 and Supplementary File 1d, e**). G-box was the most highly distributed *cis*-element present in the promoter region of the *AtSWEET11* and *AtSWEET12* orthologs from many plant species, followed by Box 4, TCT motif, GT1 motif, LAMP elements, and others. RY *cis*-elements involved in seed development were present in the promoters of a few *AtSWEET11* and *AtSWEET12* orthologs from Poaceae and a few other families (**Figure 3**). The *cis*-elements associated with phytohormone signaling were also present in the promoter sequences of *AtSWEET11* and *AtSWEET12* orthologs. The ABRE, TGACG, and CGTCA elements were highly distributed in the promoter sequences of *AtSWEET11* and *AtSWEET12* orthologs, followed by ERE, TGA, TCA, GARE elements, and others. The DRE1 element was only present in the promoter sequences of *AtSWEET12* orthologs from *Fragaria vesca* (**Figure 3**). Abiotic and biotic stress-associated *cis*-elements involved in responses to heat, cold, drought, wounding, and pathogens were also present in the promoter sequences of *AtSWEET11* and *AtSWEET12* orthologs. MYB and MYC elements were abundantly distributed in the promoter sequences of *AtSWEET11* and *AtSWEET12* orthologs from different plant species, followed by STRE, ARE, LTR, W box, WUN, and MBS elements. Defense and stress response-related TC-rich elements were only present in the promoter sequence of the *AtSWEET11* ortholog from *Solanum melongena* (**Figure 3**). Overall, *cis*-element analysis of *AtSWEET11* and *AtSWEET12* orthologs indicates that the orthologs of these transporters might play crucial roles in other plant species.

**Figure 3.**
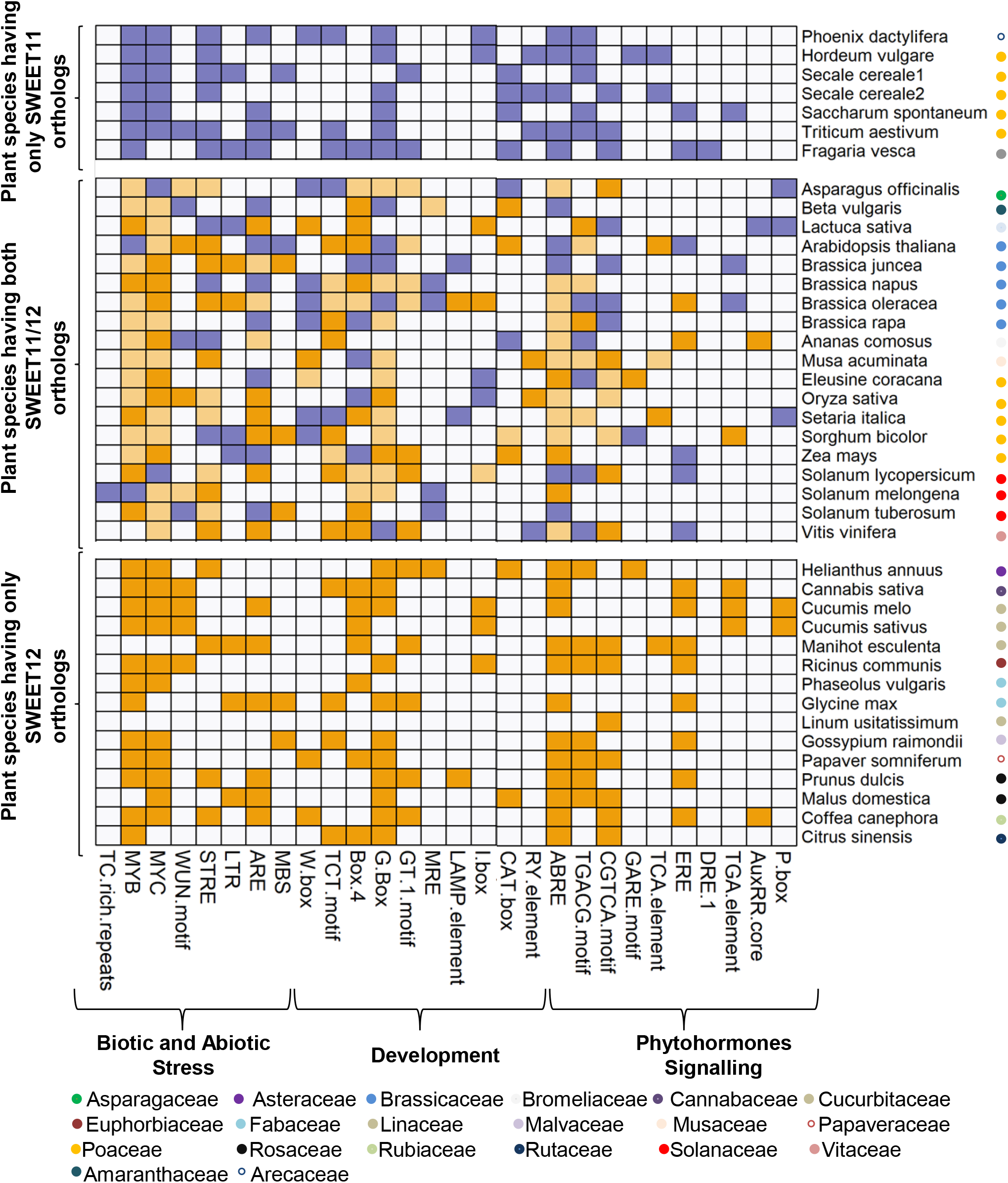
Distribution of *cis*-acting regulatory elements involved in plant development and stress responses. The *cis*-elements were identified in 1.5 kb 5′ upstream promoter regions of *AtSWEET11* and *AtSWEET12* orthologs from 39 different plant species using PlantCARE (http://bioinformatics.psb.ugent.be/webtools/plantcare/html/). The analysis involved 25 protein sequences for *AtSWEET11* orthologs and 33 protein sequences for *AtSWEET12* orthologs. The heatmap shows *cis*-elements identified in *AtSWEET11* orthologs (purple boxes), *cis*-elements identified in *AtSWEET12* orthologs (orange boxes), *cis*-elements identified in both *AtSWEET11* and *AtSWEET12* orthologs (light-orange boxes), and the absence of *cis*-elements in *AtSWEET11* and *AtSWEET12* orthologs (white boxes) from different plant species. Families of each species are indicated by colored circles. For more details, see Supplementary File 1.

Moreover, both *AtSWEET11* and *AtSWEET12* are well-expressed in the leaf tissues of Arabidopsis (Chen et al., 2012). However, in the leaves of *Oryza sativa* and *Solanum tuberosum* the transcript expression of *AtSWEET11* orthologs was high (**Supplementary Figure 8A**), while the expression of *AtSWEET12* orthologs remained undetected (**Supplementary Figure 8B**). While, in *Setaria italica* and *Sorghum bicolor*, the expression of *AtSWEET12* orthologs was high (**Supplementary Figure 8B**) and *AtSWEET11* orthologs remained undetected (**Supplementary Figure 8A**). The differential expression pattern of *AtSWEET11* and *AtSWEET12* orthologs indicate distinct and exclusive roles of these transporters in different plant species.

The length of the protein encoded by *AtSWEET11* and *AtSWEET12* orthologs varies between 209 and 302 amino acids and between 214 and 293 amino acids, respectively (**Supplementary File 1b, c**). Most AtSWEET11 and AtSWEET12 orthologs carry seven TM helices (TMH) (**Supplementary Figure 9 and Supplementary File 1a**) except two, namely, *Vitis vinifera* and *S. lycopersicum*, which form six and sixteen TMHs, respectively (**Supplementary File 1a**).

Protein motif analysis revealed the presence of 20 different motifs in both AtSWEET11 and AtSWEET12 orthologs (**Figure 4**). Of these, motifs 1, 2, 3, and 4 were identified as basic features of the transporter domain of SWEETs, while the other motifs were unannotated. Motifs 1, 2, 3, 4, and 6 were present in AtSWEET11 orthologs from all species. Motifs 1, 3, 4, 5, 6, and 9 were present in AtSWEET12 orthologs from all species (**Figure 4 and Supplementary Figure 10**). The CTD was highly non-conserved for AtSWEET11 and AtSWEET12 orthologs from different plant species (**Supplementary Figure 11**). Recently, it has been shown that phloem parenchyma cell-specific expression of AtSWEET11 is regulated at the post-transcriptional level by the two evolutionary duplicated domains in the AtSWEET11 coding sequence along with its promoter including 5′ UTR through RNA-binding protein (Zhang et al., 2021).

**Figure 4.**
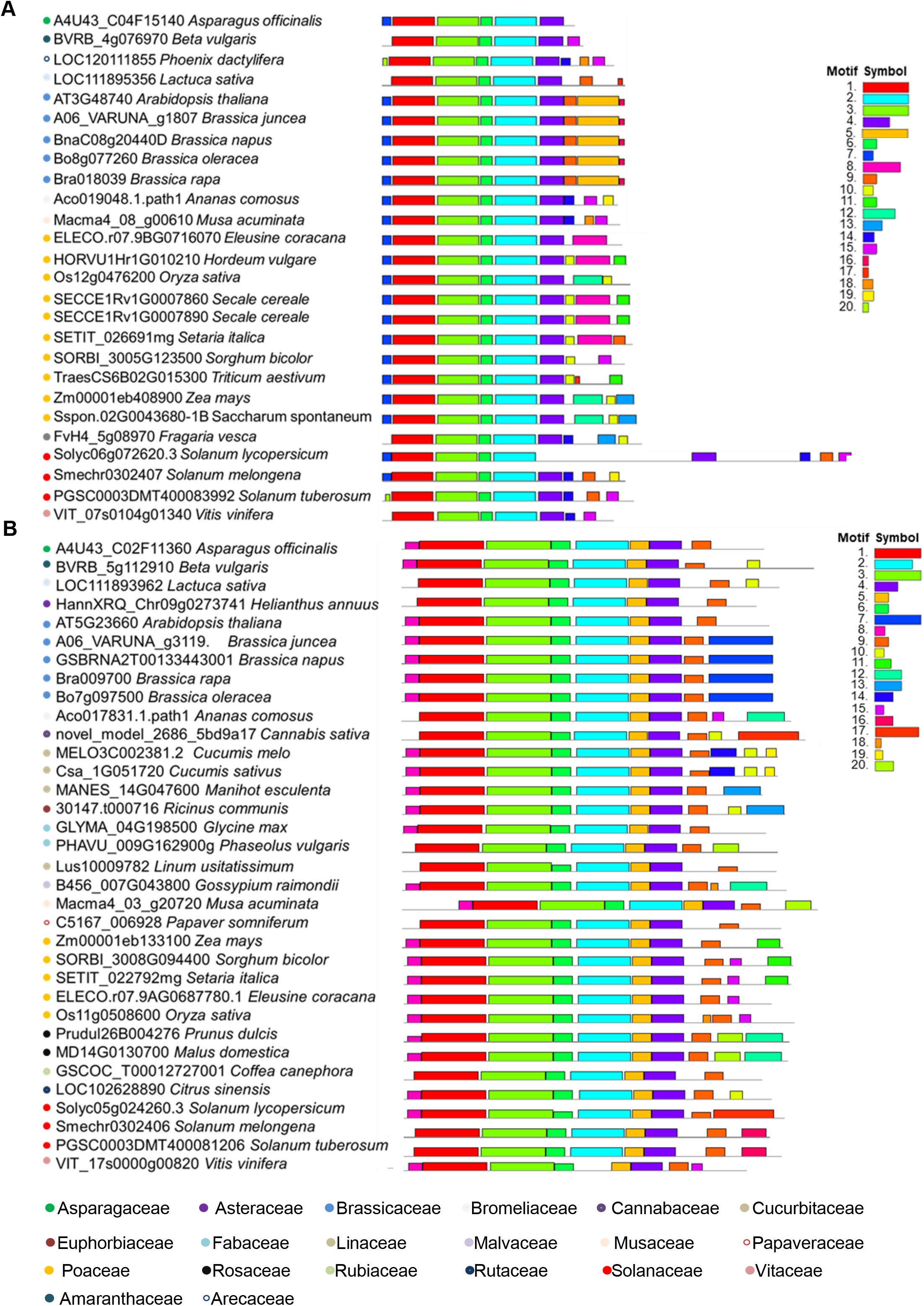
Identification of protein motifs for AtSWEET11 and AtSWEET12 orthologs in different plant species. Protein motifs were identified for the orthologs of **A**, AtSWEET11 and **B**, AtSWEET12 from 39 different plant species using MEME_suite (https://meme-suite.org/meme/). The analysis involved 25 protein sequences for A and 33 protein sequences for B. Twenty motifs were identified for each, and the motif symbols are indicated by different colors. Families of each species are indicated by colored circles. For more details, see Supplementary File 1.

Besides, transporters are mainly known to be regulated through post-translational modification (PTM) via phosphorylation, acetylation, and glycosylation. Membrane phosphoproteome analysis and the presence of a possible phosphorylation site in the CTD of AtSWEET11 (Reiland et al., 2009) strongly indicate phosphorylation-based regulation. PTM analysis of AtSWEET11 and AtSWEET12 orthologs revealed the presence of potential sites for phosphorylation and acetylation at the CTD in a majority of the analyzed plant species (Anjali et al., 2020)). However, the sites for glycosylation were available for the orthologs of only a few species (**Supplementary File 1f, g, h**). The post-translational regulation of AtSWEET11 and AtSWEET12 and their orthologs via phosphorylation and acetylation needs further experimental validation.

## Conclusion and future directions

AtSWEET11 and AtSWEET12 transporters function in tandem to modulate sugar flux in Arabidopsis. Further in-depth analysis of AtSWEET11 and AtSWEET12 transporters can provide basic molecular understanding about plant sugar transport processes. Homology models of AtSWEET11 and AtSWEET12 reveal the key amino acids essential for substrate recognition and transport. Docking studies show that the central sucrose binding residues are almost similar in both AtSWEET11 and AtSWEET12 transporters. Further *in-vitro* transport assays and *in-silico* studies might shed more light on the translocation pathway of different sugars in these transporters. Moreover, the structural similarities between AtSWEET11 and AtSWEET12 explains the functional redundancy of these two transporters. However, the CTD varies between AtSWEET11 and AtSWEET12 and their orthologs in different plant species. It is highly plausible that these transporters are independently regulated through PTM at the CTD, which might be a reason for distinct and exclusive roles of these two transporters in various plant physiological processes. Although, the phenotypic analysis for the double mutant of *AtSWEET11* and *AtSWEET12* points towards the functional redundancy (Chen et al., 2012; Gebauer et al., 2016; Walerowski et al., 2018), the histochemical and expression-based studies of the single mutants of these two transporters demonstrate their distinct roles in various developmental processes (Chen et al., 2015; Le Hir et al., 2015), as well as in response to environmental stimuli (Fatima and Senthil-Kumar, 2021). The differential expression of AtSWEET11 and AtSWEET12 orthologs also support their specific roles in different plant species (**Supplementary Figure 8**). Moreover, it has been shown that the phloem-specific expression of *AtSWEET11* is controlled at post-transcriptional level (Zhang et al., 2021). More studies are needed to understand the regulation of these two transporters in controlling sugar availability to pathogens and in restricting disease development. At post-translational level, oligomerization and PTM could be likely mechanisms of regulation for AtSWEET11 and AtSWEET12 transporters. The biochemical transport assays suggest an oligomerization-based function of SWEETs (Xuan et al., 2013; Han et al., 2017). AtSWEET11 and AtSWEET12 could homo- or heterodimerize with each other (Xuan et al., 2013; Fatima and Senthil-Kumar, 2021) and the hetero-oligomerization of AtSWEET11 and AtSWEET12 impacts the sucrose transport activity (Fatima and Senthil-Kumar, 2021).

Although research on SWEETs has increased considerably, the mechanism of regulation of SWEETs in crop plants during various developmental stages and abiotic and biotic stress conditions remains to be elucidated in greater detail. The structure– function relationship of AtSWEET11 and AtSWEET12 deciphered from the experimentally determined structures with their corresponding native sugar substrates would help gain deeper insights into the mechanistic details of the transport. Finer details on the orthologs will help implement experimental concepts from model plant to crop plants to develop improved crop phenotypes with enhanced productivity and resilience.

## Methodology

### Molecular dynamics (MD) trajectory analysis

The full-length amino acid sequences of AtSWEET11 and AtSWEET12 were obtained from TAIR (https://www.arabidopsis.org) and submitted to psiPRED server (http://bioinf.cs.ucl.ac.uk/psipred) for sequence based secondary structure prediction. Psipred predicted transmembrane helices and cytoplasmic disordered C-terminal region for both the sequences. For homology modelling, the full length AtSWEET11 and AtSWEET12 protein sequence were submitted to the Robetta server (http://robetta.bakerlab.org) and the output models for AtSWEET11 (PD algorithm) and AtSWEET12 (TR algorithm) with 0.76 and 0.74 confidence score respectively were selected after visualising them. The models selected had minimal estimated positional error for the residues. However, the C-terminal for both the predicted proteins was highly disordered with maximal regions modelled as loops. The 3D coordinates of AtSWEET11 and AtSWEET12 homology models were prepared using the protein preparation wizard of the Schrodinger suite. The missing residues and side chains were filled and bond order errors were corrected, followed by H-bond optimisation and a restrained minimisation with a cut-off of 0.3 Å. The prepared protein models were simulated to equilibrate in a POPC lipid bilayer at 300 K. POPC lipid was automatically placed perpendicular to the transmembrane helices and the systems were solvated using TIP3P water model. The MD system was electrostatically neutralised by placing counter Cl^-^ ions and 0.15 M KCl was added to provide adequate ionic strength. The prepared systems were relaxed by default desmond relaxation protocols before being simulated for 600ns with a recording interval of 50ps. The RMSD plots of the simulated MD runs indicated that the transmembrane region stabilised with an average c-alpha RMSD of 2.3 Å and 2.06 Å for AtSWEET11 and AtSWEET12 respectively. The C-terminal domain was found to be highly disordered as reflected by high RMSF values and contributed majorly to the overall RMSD of the whole structure throughout the simulation.

### RMSD plot

The 3D coordinates of sucrose were extracted from PDB:3LDK and possible conformers were generated using Ligprep. The 3D coordinates of the AtSWEET11 and AtSWEET12 transmembrane domain were extracted from the last frame of the 600ns MD and subjected to a sitemap analysis. The best scoring sites with site scores of 1.139 and 1.134 were predicted at the central cavity for AtSWEET11 and AtSWEET12 respectively. These sites were selected to generate receptor grids for docking sucrose. The sucrose was docked at the prepared grid using the XP (Extra precision) docking mode using Glide. Output poses were visualised for interactions and clashes, and binding energy calculations were calculated through Prime-MMGBSA module of Schrodinger suite to select the best pose for MD run. The sucrose docked AtSWEET11 and AtSWEET12 were prepared in lipid bilayer membrane environment and simulated for 500ns and 1us respectively to study their interaction dynamics. Thermal MM-GBSA script was ran on the MD trajectory of AtSWEET11 and AtSWEET12 to obtain dG binding score w.r.t. Frames. The most stable complex according to the dG binding score was exported to study interactions/representation. The MD trajectory analysis for AtSWEET11 and AtSWEET12 sucrose docked poses revealed that the c-alphas were stable throughout the simulation with an average RMSD of 1.43 Å and 1.93 Å respectively, while RMSD for sucrose was calculated to be 4.42 Å and 2.33 Å respectively.

### Tertiary structure prediction

The tertiary structures of the proteins orthologus to SWEET11 and SWEET12 were predicted using Robetta (https://robetta.bakerlab.org/).

### Multiple sequence alignment

The full-length amino acid sequences of AtSWEET11, AtSWEET12, AtSWEET13 and OsSWEET2b were obtained from TAIR (https://www.arabidopsis.org) Sequences were aligned using ClustalW (https://www.genome.jp/tools-bin/clustalw).

### Time-tree analysis

The phylogenetic analysis of thirty-nine different plant species from twenty different families was performed. The species names were taken as input and time-tree was generated using the MEGA X (1). The nodes in the tree indicate the divergence times for different plant species.

### Gene structure analysis

The gene structure analysis including exon-intron arrangement were conducted for the orthologs of AtSWEET11 gene and AtSWEET12 gene from thirty-nine different plant species using Gene Structure Display Server 2.0 (http://gsds.gao-lab.org/index.php). The analysis involved 25 gene sequences for 11, and 33 gene sequences for 12.

### Phylogenetic analysis

The evolutionary analysis of AtSWEET11 and AtSWEET12 orthologs in thirty-nine different plant species from twenty different families was performed. The time-tree was generated using the RelTime method.

### Protein motif analysis

The logo of each motif and associated amino acids were identified for AtSWEET11 and AtSWEET12 protein orthologs from different plant species using MEME_suite (https://meme-suite.org/meme/).

## Acknowledgements

The SWEET transporter project at MS-K’s lab was funded by NIPGR core funding. UF acknowledges the DBT-SRF fellowship (DBT/2013/NIPGR/68) and NIPGR-SRF fellowship. DB is supported by CSIR-SRA fellowship award and WAK is supported by DBT fellowship. Authors acknowledge the use of Schrodinger suite licensed to ICGEB. AA lab is supported by ICGEB core funds. Authors acknowledge Dr. Aashish Ranjan and Miss Anjali for critical reading of the manuscript.

## Author contribution statement

MS-K conceived the concept and provided outline. UF contributed to the figures, supplemental information and drafted the entire manuscript. DB, WAK and AA contributed to the structure analysis and related part of the manuscript. MK contributed to motif analysis, cis element analysis and related supplementary figures. JV involved in editing the manuscript. MS-K edited and finalized the manuscript.

## List of supplements

Supplementary Figure 1. The Molecular dynamics (MD) trajectory analysis for AtSWEET11 and AtSWEET12

Supplementary Figure 2. Structural superposition of AtSWEET11 and AtSWEET12.

Supplementary Figure 3. RMSD plot of AtSWEET11 and AtSWEET12 for protein backbone and sucrose as a ligand.

Supplementary Figure 4. Multiple sequence alignment of AtSWEET11, AtSWEET12, AtSWEET13, and OsSWEET2b.

Supplementary Figure 5. The time-tree analysis for evaluating the divergence time of different species used in this study.

Supplementary Figure 6. Phylogenetic analysis of AtSWEET11 and AtSWEET12 orthologs in different plant species.

Supplementary Figure 7. Exon–intron distribution of AtSWEET11 and AtSWEET12 orthologs in different plant species.

Supplementary Figure 8. Transcript expression of AtSWEET11 and AtSWEET12 orthologs in the leaf tissues from different plant species.

Supplementary Figure 9. Tertiary structure prediction of AtSWEET11 and AtSWEE12 orthologs from different plant species.

Supplementary Figure 10. Details of logos of each protein motif for AtSWEET11 and AtSWEET12 orthologs in different plant species.

Supplementary Figure 11. The C-terminal analysis of AtSWEET11 and AtSWEET12 protein orthologs from different plant species.

Supplementary Table 1: Sucrose-interacting residues in AtSWEET11 and AtSWEET12 with equivalent dCMP-binding residues in the AtSWEET13 crystal structure.

Supplementary Table 2: Details of conserved residues among AtSWEETs and OsSWEET2b.

Supplementary File S1. Details regarding data used for figure preparation.

## References

Anjali A., Fatima U., Manu M.S., Ramasamy S., Senthil-Kumar M. Structure and regulation of SWEET transporters in plants: An update. Plant Phys. Biochem. 2020; 156: 1–6

Antony G., Zhou J., Huang S., Li T., Liu B., White F., Yang B. Rice xa13 recessive resistance to bacterial blight is defeated by induction of the disease susceptibility gene Os-11N3. Plant Cell. 2010; 22: 3864–3876.

Ayre B.G. Membrane-transport systems for sucrose in relation to whole-plant carbon partitioning. Mol. Plant. 2011; 4: 377–394.

Breia R., Conde A., Badim H., Fortes A.M., Gerós H., Granell A. Plant SWEETs: from sugar transport to plant–pathogen interaction and more unexpected physiological roles. Plant Physiol. 2021; 186: 836–852

Chen L.Q., Hou B.H., Lalonde S., Takanaga H., Hartung M.L., Qu X.Q., Guo W.J., Kim J.G., Underwood W., Chaudhuri B., Chermak D. Sugar transporters for intercellular exchange and nutrition of pathogens. Nature. 2010; 468: 527–532.

Chen L.Q., Qu X.Q., Hou B.H., Sosso D., Osorio S., Fernie A.R., Frommer W.B. Sucrose efflux mediated by SWEET proteins as a key step for phloem transport. Science. 2012; 335: 207–211.

Chen L.Q., Lin I.W., Qu X.Q., Sosso D., McFarlane H.E., Londoño A., Samuels A.L., Frommer W.B. A cascade of sequentially expressed sucrose transporters in the seed coat and endosperm provides nutrition for the Arabidopsis embryo. Plant Cell. 2015; 27: 607–619.

Chen, Q., Hu, T., Li, X., Song, C. P., Zhu, J. K., Chen, L., Zhao, Y. Phosphorylation of SWEET sucrose transporters regulates plant root:shoot ratio under drought. Nat. Plants2022; 8: 68–77.

Chong J., Piron M.C., Meyer S., Merdinoglu D., Bertsch C., Mestre P. The SWEET family of sugar transporters in grapevine: VvSWEET4 is involved in the interaction with Botrytis cinerea. J. Exp. Bot. 2014; 65: 6589–6601.

Cox K.L., Meng F., Wilkins K.E., Li F., Wang P., Booher N.J. TAL effector driven induction of a SWEET gene confers susceptibility to bacterial blight of cotton. Nature Commun. 2017; 8: 1–14.

Durand M., Porcheron B., Hennion N., Maurousset L., Lemoine R., Pourtau N. Water deficit enhances C export to the roots in Arabidopsis thaliana plants with contribution of sucrose transporters in both shoot and roots. Plant Physiol. 2016; 170: 1460–1479.

Desrut A., Moumen B., Thibault F., Le Hir R., Coutos-Thévenot P. Beneficial rhizobacteria Pseudomonas simiae WCS417 induce major transcriptional changes in plant sugar transport. J. Exp. Botany. 2020; 71: 7301–7315

Fatima U., Senthil-Kumar M. Sweet revenge: AtSWEET12 in plant defense against bacterial pathogens by apoplastic sucrose limitation. bioRxiv. 2021 https://www.biorxiv.org/content/10.1101/2021.10.04.463061v1

Feng L., Frommer W.B. Structure and function of SemiSWEET and SWEET sugar transporters. Trends Biochem. Sci. 2015; 40: 480–486.

Gebauer P., Korn M., Engelsdorf T., Sonnewald U., Koch C., Voll L.M. Sugar accumulation in leaves of Arabidopsis sweet11/sweet12 double mutants enhances priming of the salicylic acid-mediated defense response. Front. Plant Sci. 2017; 8:1378.

Giaquinta R.T. Phloem loading of sucrose. Annu. Rev. Plant Physiol. Plant Mol. Bio. 1983; 34: 347–387.

Gong, Z., Yang, S. Drought meets SWEET. Nat. Plants 2022; 8: 25–26.

Rouina H., Tseng Y.H., Nataraja K.N., Uma Shaanker R., Oelmüller R. Arabidopsis restricts sugar loss to a colonizing Trichoderma harzianum strain by downregulating SWEET11 and-12 and upregulation of SUC1 and SWEET2 in the roots. Microorganisms. 2021 9: 1246.

Han L., Zhu Y., Liu M., Zhou Y., Lu G., Lan L., Wang X., Zhao Y., Zhang X.C. Molecular mechanism of substrate recognition and transport by the AtSWEET13 sugar transporter. Proc. Natl. Acad. Sci. U S A. 2017; 114: 10089–10094.

Jumper, J., Evans, R., Pritzel, A., Green, T., Figurnov, M., Ronneberger, O., Tunyasuvunakool, K., Bates, R., Žídek, A., Potapenko, A. and Bridgland, A. Highly accurate protein structure prediction with AlphaFold. Nature 2021; 596, 583–589.

Li T., Liu B., Spalding M.H., Weeks D.P., Yang B. High-efficiency TALEN-based gene editing produces disease-resistant rice. Nature Biotech. 2012; 30: 390–392.

Li H., Li X., Xuan Y., Jiang J., Wei Y., Piao Z. Genome wide identification and expression profiling of SWEET genes family reveals its role during Plasmodiophora brassicae-induced formation of clubroot in Brassica rapa. Front. Plant Sci. 2018; 9: 207.

Le Hir R., Spinner L., Klemens P.A., Chakraborti D., de Marco F., Vilaine F., Wolff N., Lemoine R., Porcheron B., Géry C., Téoulé E. Disruption of the sugar transporters AtSWEET11 and AtSWEET12 affects vascular development and freezing tolerance in Arabidopsis. Mol. Plant. 2015; 8: 1687–1690.

Papaioannou et al. SWEET genes mediate sugar translocation and allocation as a key step in roots of in vitro grown Arabidopsis. Thesis, 2018; https://edepot.wur.nl/460386

Reiland S., Messerli G., Baerenfaller K., Gerrits B., Endler A., Grossmann J., Gruissem W., Baginsky S. Large-scale Arabidopsis phosphoproteome profiling reveals novel chloroplast kinase substrates and phosphorylation networks. Plant Physiol. 2009; 150: 889–903.

Ruan Y.L., Jin Y., Yang Y.J., Li G.J., Boyer J.S. Sugar input, metabolism, and signaling mediated by invertase: roles in development, yield potential, and response to drought and heat. Mol. Plant. 2010; 3: 942–955.

Srivastava A.C., Ganesan S., Ismail I.O., Ayre B.G. Functional characterization of the Arabidopsis AtSUC2 sucrose/H+ symporter by tissue-specific complementation reveals an essential role in phloem loading but not in long-distance transport. Plant Physiol. 2008; 148: 200–211.

Srivastava A.C., Dasgupta K., Ajieren E., Costilla G., McGarry R.C., Ayre B.G. Arabidopsis plants harbouring a mutation in AtSUC2, encoding the predominant sucrose/proton symporter necessary for efficient phloem transport, are able to complete their life cycle and produce viable seed. Ann. Botany. 2009a; 104: 1121–1128.

Srivastava A., Ganesan S., Ismail I., Ayre B. Effective carbon partitioning driven by exotic phloem-specific regulatory elements fused to the Arabidopsis thaliana AtSUC2 sucrose-proton symporter gene. BMC Plant Bio. 2009b; 9: 7.

Stadler R., Lauterbach C., Sauer N. Cell-to-cell movement of green fluorescent protein reveals post-phloem transport in the outer integument and identifies symplastic domains in Arabidopsis seeds and embryos. Plant Physiol. 2005; 139: 701–712.

Tao Y., Cheung L.S., Li S., Eom J.S., Chen L.Q., Xu Y., Perry K., Frommer W.B., Feng L. Structure of a eukaryotic SWEET transporter in a homotrimeric complex. Nature. 2015; 527: 259–263.

Varadi, M., Anyango, S., Deshpande, M., Nair, S., Natassia, C., Yordanova, G., Yuan, D., Stroe, O., Wood, G., Laydon, A. and Žídek, A. AlphaFold Protein Structure Database: Massively expanding the structural coverage of protein-sequence space with high-accuracy models. Nucleic Acids Research, 2022; 50: D439–D444.

Walerowski P., Gündel A., Yahaya N., Truman W., Sobczak M., Olszak M., Rolfe S., Borisjuk L., Malinowski R. Clubroot disease stimulates early steps of phloem differentiation and recruits SWEET sucrose transporters within developing galls. Plant Cell. 2018; 30: 3058–3073.

Wei X., Nguyen S.T., Collings D.A., McCurdy D.W. Sucrose regulates wall ingrowth deposition in phloem parenchyma transfer cells in Arabidopsis via affecting phloem loading activity. J. Exp. Botany. 2020; 71: 4690–4702.

Xuan Y.H., Hu Y.B., Chen L.Q., Sosso D., Ducat D.C., Hou B.H., Frommer W.B. Functional role of oligomerization for bacterial and plant SWEET sugar transporter family. Proc. Natl. Acad. Sci U S A. 2013; 110: E3685–3694.

Yang J., Anishchenko I., Park H., Peng Z., Ovchinnikov S., Baker D. Improved protein structure prediction using predicted interresidue orientations. Proc. Natl. Acad. Sci. U S A. 2020; 117: 1496–1503.

Zhang C., Li Y., Wang J., Xue X., Beuchat G., Chen LQ. Two evolutionarily duplicated domains individually and post-transcriptionally control SWEET expression for phloem transport. New Phytol. 2021; 10.1111/nph.17688.

