## Supplemental information (figures and tables) for "SWEET11 and SWEET12 transporters function in tandem to modulate sugar flux in Arabidopsis: An account of the underlying unique structure–function relationship"

### Supplementary figures and their legends

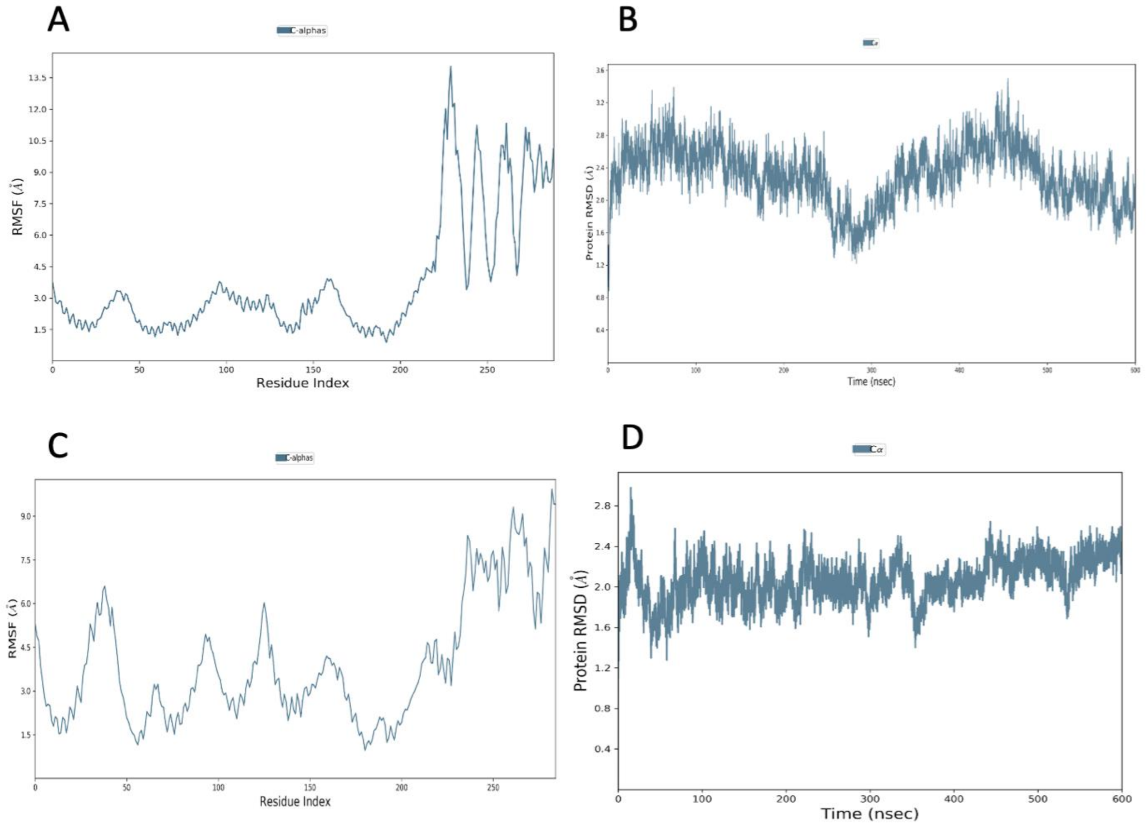

**Supplementary Figure 1. The Molecular dynamics (MD) trajectory analysis for AtSWEET11 and AtSWEET12 A,** RMSF plot for full length AtSWEET11. **B,** RMSD plot for the TM region of AtSWEET11 **C,** RMSF plot for full length AtSWEET12 **D,** RMSD plot for the TM region of AtSWEET12.

The full-length amino acid sequences of AtSWEET11 and AtSWEET12 were obtained from TAIR (https://www.arabidopsis.org) and submitted to psiPRED server (http://bioinf.cs.ucl.ac.uk/psipred) for sequence based secondary structure prediction. Psipred predicted transmembrane helices and cytoplasmic disordered C-terminal region for both the sequences. For homology modelling, the full length AtSWEET11 and AtSWEET12 protein sequence were submitted to the Robetta server (http://robetta.bakerlab.org) and the output models for AtSWEET11 (PD algorithm) and AtSWEET12 (TR algorithm) with 0.76 and 0.74 confidence score respectively were selected after visualising them. The models selected had minimal estimated positional error for the residues. However, the C-terminal for both the predicted proteins was highly disordered with maximal regions modelled as loops. The 3D coordinates of AtSWEET11 and AtSWEET12 homology models were prepared using the protein preparation wizard of the Schrodinger suite. The missing residues and side chains were filled and bond order errors were corrected, followed by H-bond optimisation and a restrained minimisation with a cut-off of 0.3 Å. The prepared protein models were simulated to equilibrate in a POPC lipid bilayer at 300 K. POPC lipid was automatically placed perpendicular to the transmembrane helices and the systems were solvated using TIP3P water model. The MD system was electrostatically neutralised by placing counter $Cl^-$ ions and 0.15 M KCl was added to provide adequate ionic strength. The prepared systems were relaxed by default desmond relaxation protocols before being simulated for 600ns with a recording interval of 50ps. The RMSD plots of the simulated MD runs indicated that the transmembrane region stabilised with an average c-alpha RMSD of 2.3 Å and 2.06 Å for AtSWEET11 and AtSWEET12 respectively. The C-terminal domain was found to be highly disordered as reflected by high RMSF values and contributed majorly to the overall RMSD of the whole structure throughout the simulation. The C-terminal domain was excluded and only the stable transmembrane domain of 1-219 amino acids was used for docking and MD analysis.

Sastry, G.M.; Adzhigirey, M.; Day, T.; Annabhimoju, R.; Sherman, W., "Protein and ligand preparation: Parameters, protocols, and influence on virtual screening enrichments," J. Comput. Aid. Mol. Des., 2013, 27(3), 221-234

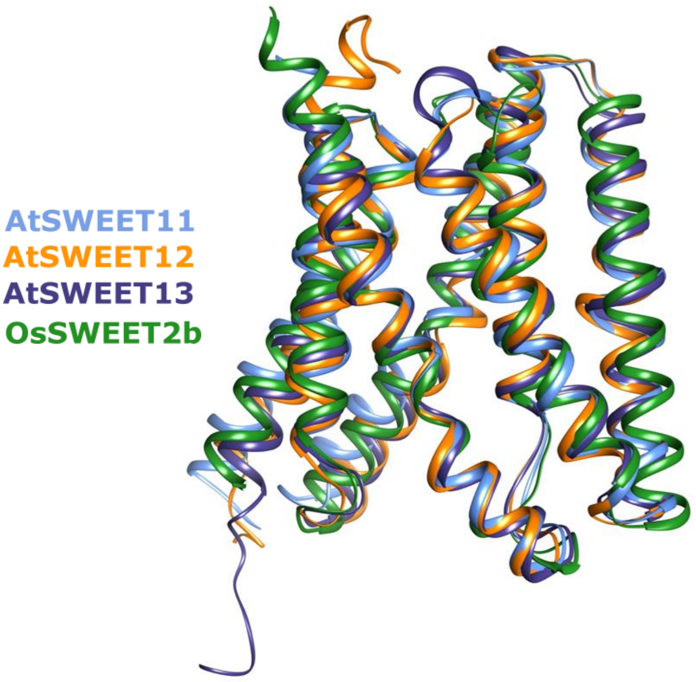

**Supplementary Figure 2. Structural superposition of AtSWEET11, AtSWEET12, AtSWEET13 and OsSWEET2b.** Superposition of homology models of AtSWEET11 (cornflower blue) and AtSWEET12 (orange) with crystal structures of AtSWEET13 (violet) and OsSWEET2b (forest green). The RMSD values of superposition of AtSWEET11 with AtSWEET13 and OsSWEET2b is 1.66 Å and 2.56 Å respectively. The RMSD values of superposition of AtSWEET12 with AtSWEET13 and OsSWEET2b is 1.36 Å and 2.50 Å respectively. For structural superposition, AtSWEET13 (PDB: 5XPD Chain A 1-222 residues), OsSWEET2b (PDB: 5CTG Chain A 1-215 residues) and homology models of AtSWEET11 and AtSWEET12 (1-219 residues) were used. RMSD values were calculated through SSM superposition in COOT. For more details see Supplementary Figure 1

A B

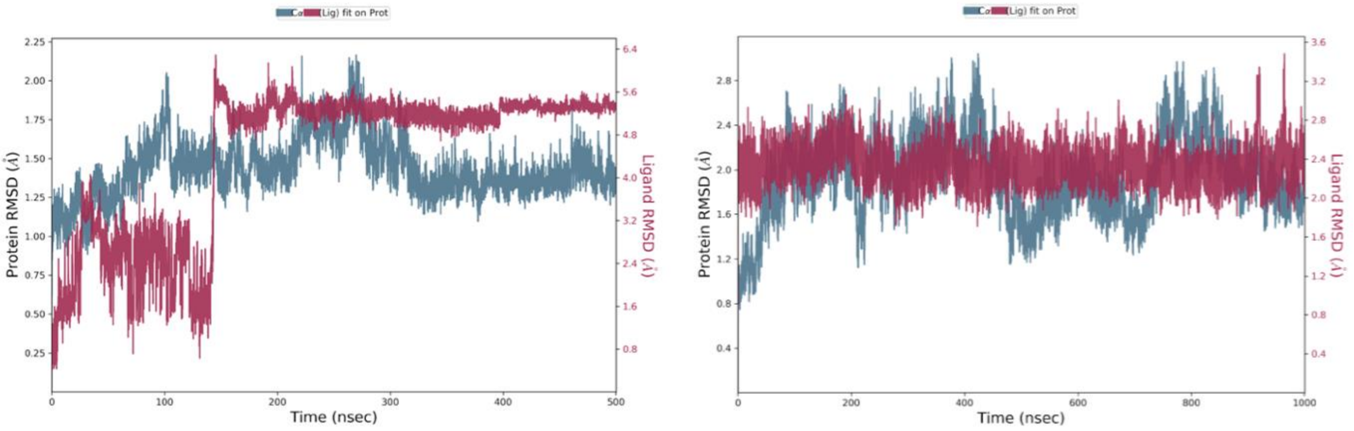

**Supplementary Figure 3. The RMSD plot of AtSWEET11 and AtSWEET12 for protein backbone and sucrose as ligand. A,** The RMSD plot of AtSWEET11. **B,** The RMSD plot of AtSWEET12. The 3D coordinates of sucrose were extracted from PDB:3LDK and possible conformers were generated using Ligprep. The 3D coordinates of the AtSWEET11 and AtSWEET12 transmembrane domain were extracted from the last frame of the 600ns MD and subjected to a sitemap analysis. The best scoring sites with site scores of 1.139 and 1.134 were predicted at the central cavity for AtSWEET11 and AtSWEET12 respectively. These sites were selected to generate receptor grids for docking sucrose. The sucrose was docked at the prepared grid using the XP (Extra precision) docking mode using Glide. Output poses were visualised for interactions and clashes, and binding energy calculations were calculated through Prime-MMGBSA module of Schrodinger suite to select the best pose for MD run. The sucrose docked AtSWEET11 and AtSWEET12 were prepared in lipid bilayer membrane environment and simulated for 500ns and 1us respectively to study their interaction dynamics. Thermal MM-GBSA script was ran on the MD trajectory of AtSWEET11 and AtSWEET12 to obtain dG binding score w.r.t. Frames. The most stable complex according to the dG binding score was exported to study interactions/representation. The MD trajectory analysis for AtSWEET11 and AtSWEET12 sucrose docked poses revealed that the c-alphas were stable throughout the simulation with an average RMSD of 1.43 Å and 1.93 Å respectively, while RMSD for sucrose was calculated to be 4.42 Å and 2.33 Å respectively.

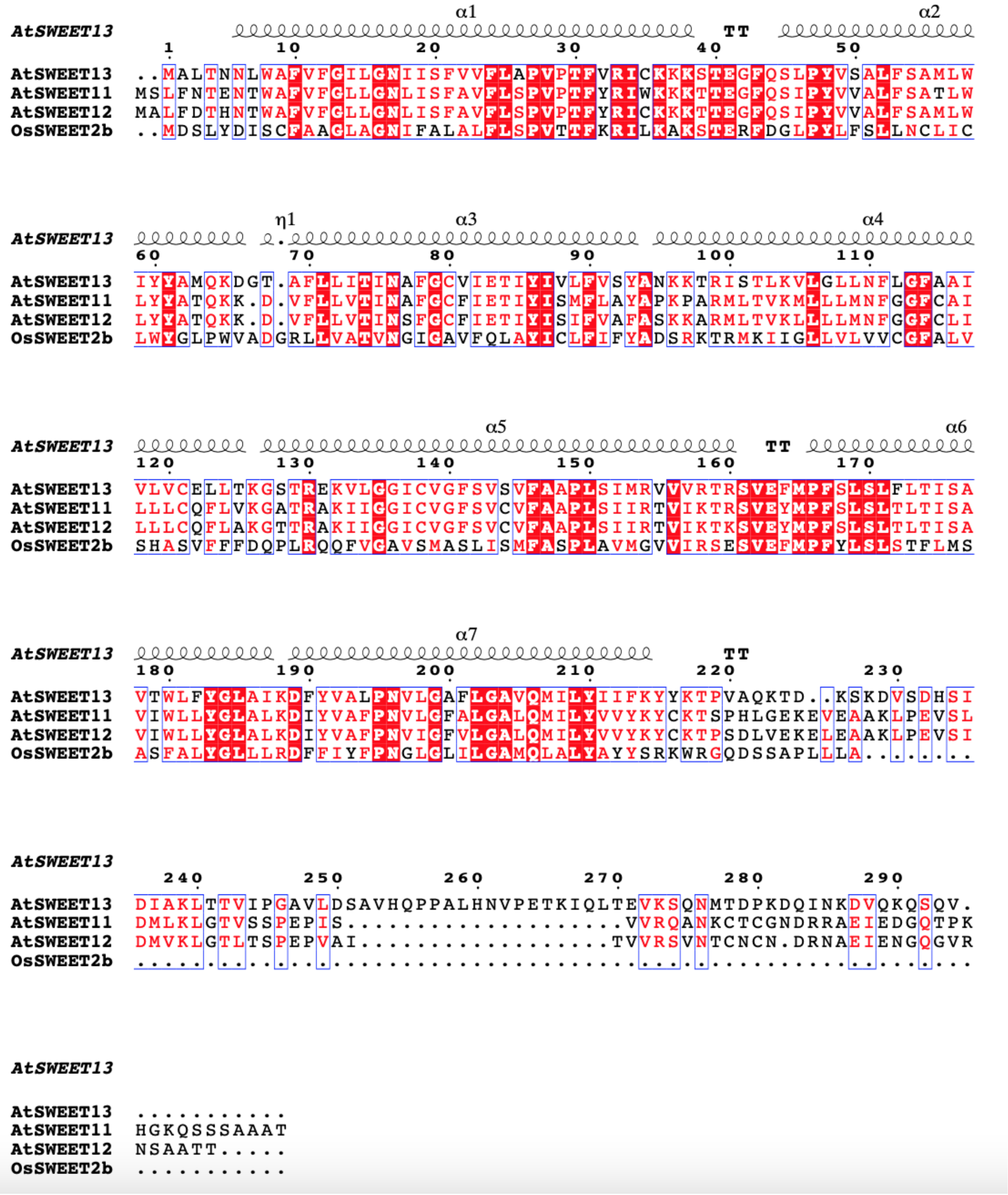

**Supplementary Figure 4. Multiple sequence alignment of AtSWEET11, AtSWEET12, AtSWEET13 and OsSWEET2b.**

The full-length amino acid sequences of AtSWEET11, AtSWEET12, AtSWEET13 and OsSWEET2b were obtained from TAIR (https://www.arabidopsis.org) Sequences were aligned using ClustalW (https://www.genome.jp/tools-bin/clustalw). Secondary structure assignments of AtSWEET13 are indicated above the alignment. Multiple sequence alignment of AtSWEET11, AtSWEET12 with AtSWEET13 and OsSWEET2b showed a total of 61 conserved residues. The conserved residues are highlighted in red.

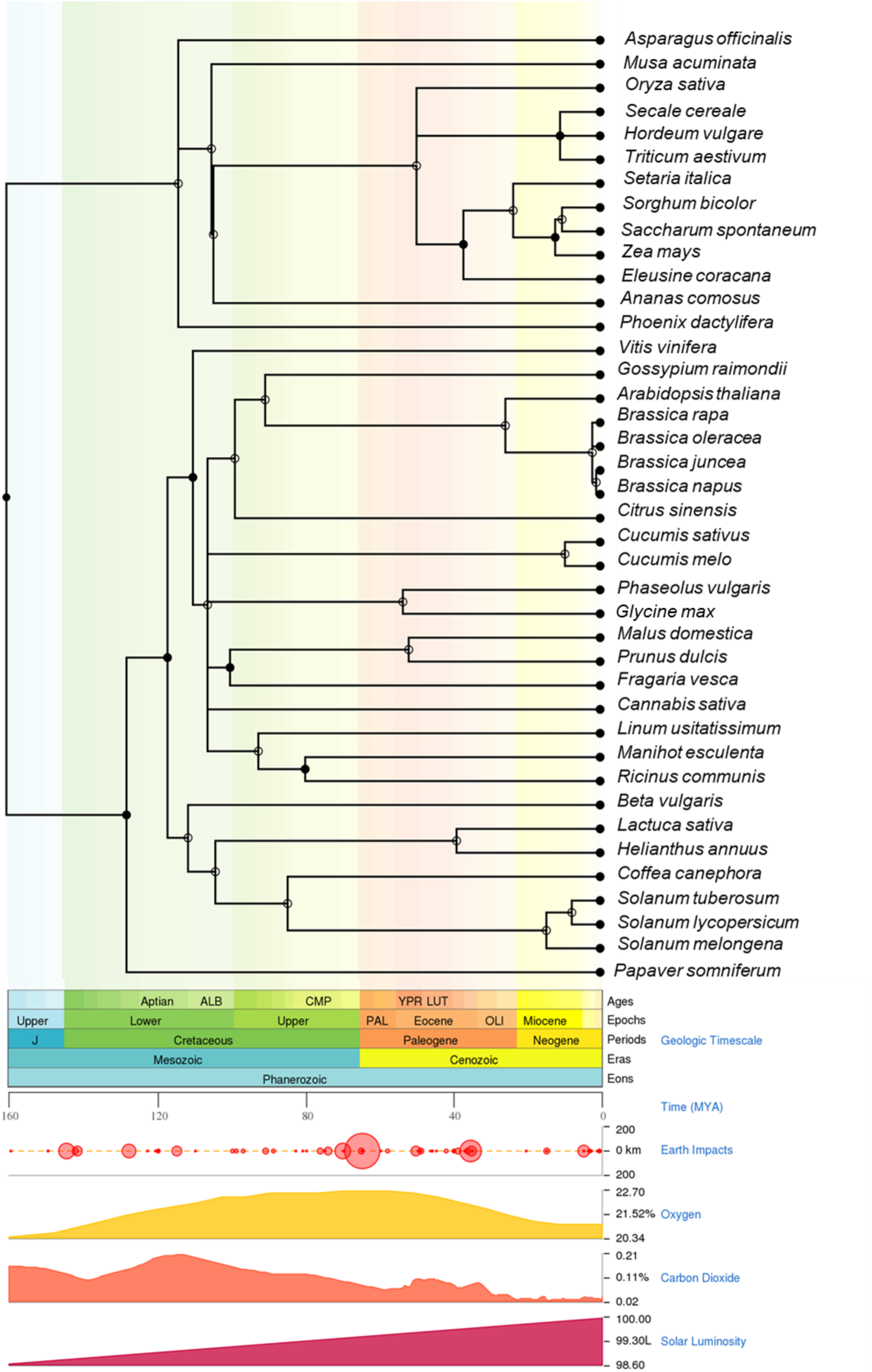

Asparagus officinalis
Musa acuminata
Oryza sativa
Secale cereale
Hordeum vulgare
Triticum aestivum
Setaria italica
Sorghum bicolor
Saccharum spontaneum
Zea mays
Eleusine coracana
Ananas comosus
Phoenix dactylifera
Vitis vinifera
Gossypium raimondii
Arabidopsis thaliana
Brassica rapa
Brassica oleracea
Brassica juncea
Brassica napus
Citrus sinensis
Cucumis sativus
Cucumis melo
Phaseolus vulgaris
Glycine max
Malus domestica
Prunus dulcis
Fragaria vesca
Cannabis sativa
Linum usitatissimum
Manihot esculenta
Ricinus communis
Beta vulgaris
Lactuca sativa
Helianthus annuus
Coffea canephora
Solanum tuberosum
Solanum lycopersicum
Solanum melongena
Papaver somniferum
Aptian
ALB
CMP
YPR LUT
Ages
Upper
Lower
Upper
PAL
Eocene
OLI
Miocene
Epochs
J
Cretaceous
Paleogene
Neogene
Periods
Geologic Timescale
Mesozoic
Cenozoic
Eras
Phanerozoic
Eons
160
120
80
40
0
Time (MYA)
200
0 km
200
Earth Impacts
22.70
21.52%
20.34
Oxygen
0.21
0.11%
0.02
Carbon Dioxide
100.00
99.30L
98.60
Solar Luminosity

**Supplementary Figure 5. The time-tree analysis for evaluating the divergence time of different species used in this study.**

The phylogenetic analysis of thirty-nine different plant species from twenty different families was performed. The species names were taken as input and  time-tree was generated using the MEGA X (1). The nodes in the tree indicate the divergence times for different plant species. The complete list of the species can obtained from Supplementary File 1.

Kumar S., Stecher G., Li M., Knyaz C., and Tamura K. (**2018**). MEGA X: Molecular Evolutionary Genetics Analysis across computing platforms. *Molecular Biology and Evolution* **35**:1547-1549.

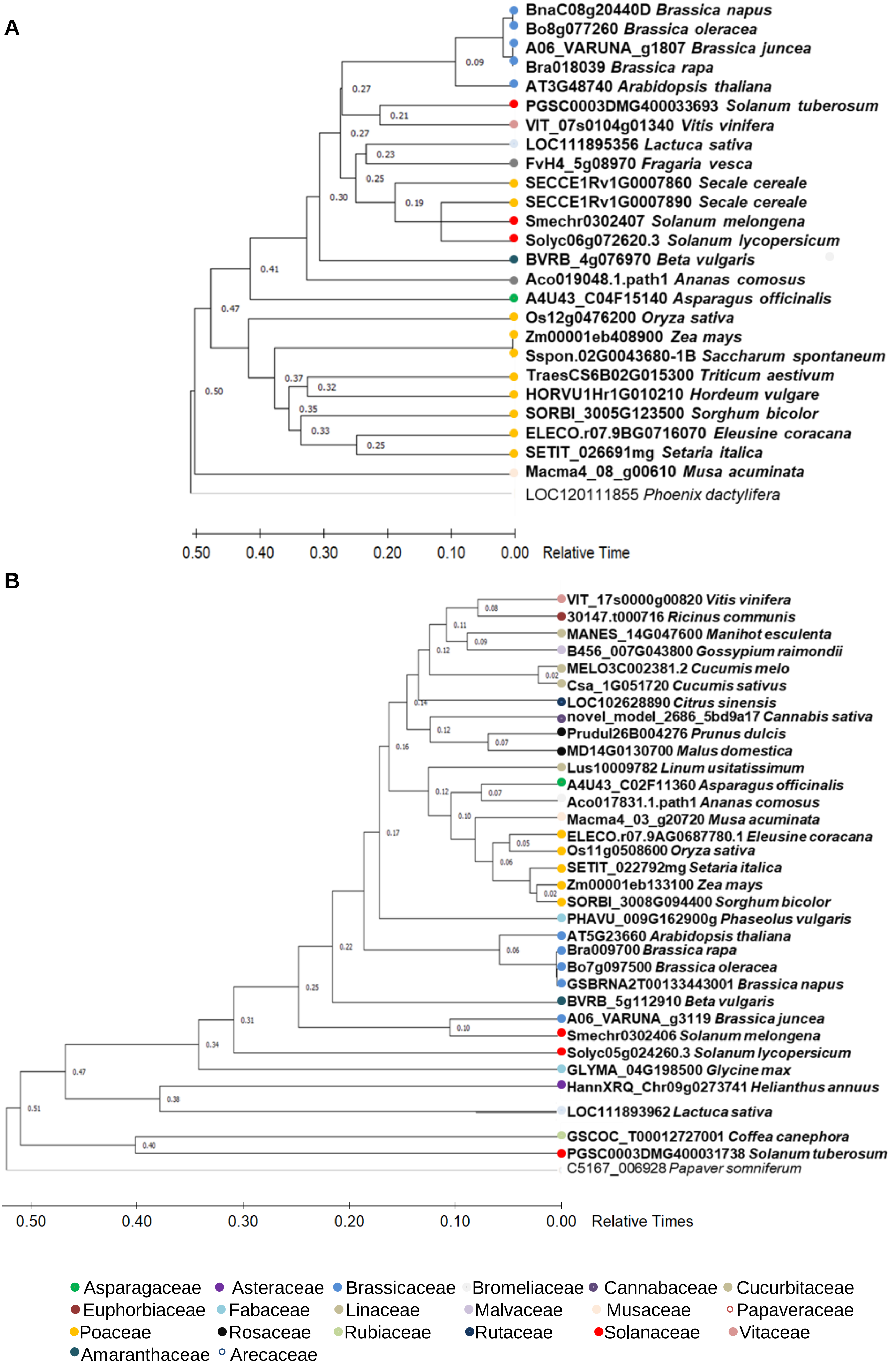
A
BnaC08g20440D Brassica napus
Bo8g077260 Brassica oleracea
A06_VARUNA_g1807 Brassica juncea
Bra018039 Brassica rapa
AT3G48740 Arabidopsis thaliana
PGSC0003DMG400033693 Solanum tuberosum
VIT_07s0104g01340 Vitis vinifera
LOC111895356 Lactuca sativa
FvH4_5g08970 Fragaria vesca
SECCE1Rv1G0007860 Secale cereale
SECCE1Rv1G0007890 Secale cereale
Smechr0302407 Solanum melongena
Solyc06g072620.3 Solanum lycopersicum
BVRB_4g076970 Beta vulgaris
Aco019048.1.path1 Ananas comosus
A4U43_C04F15140 Asparagus officinalis
Os12g0476200 Oryza sativa
Zm00001eb408900 Zea mays
Sspon.02G0043680-1B Saccharum spontaneum
TraesCS6B02G015300 Triticum aestivum
HORVU1Hr1G010210 Hordeum vulgare
SORBI_3005G123500 Sorghum bicolor
ELECO.r07.9BG0716070 Eleusine coracana
SETIT_026691mg Setaria italica
Macma4_08_g00610 Musa acuminata
LOC120111855 Phoenix dactylifera
0.50 0.40 0.30 0.20 0.10 0.00 Relative Time
B
VIT_17s0000g00820 Vitis vinifera
30147.t000716 Ricinus communis
MANES_14G047600 Manihot esculenta
B456_007G043800 Gossypium raimondii
MELO3C002381.2 Cucumis melo
Csa_1G051720 Cucumis sativus
LOC102628890 Citrus sinensis
novel_model_2686_5bd9a17 Cannabis sativa
Prudul26B004276 Prunus dulcis
MD14G0130700 Malus domestica
Lus10009782 Linum usitatissimum
A4U43_C02F11360 Asparagus officinalis
Aco017831.1.path1 Ananas comosus
Macma4_03_g20720 Musa acuminata
ELECO.r07.9AG0687780.1 Eleusine coracana
Os11g0508600 Oryza sativa
SETIT_022792mg Setaria italica
Zm00001eb133100 Zea mays
SORBI_3008G094400 Sorghum bicolor
PHAVU_009G162900g Phaseolus vulgaris
AT5G23660 Arabidopsis thaliana
Bra009700 Brassica rapa
Bo7g097500 Brassica oleracea
GSBRNA2T00133443001 Brassica napus
BVRB_5g112910 Beta vulgaris
A06_VARUNA_g3119 Brassica juncea
Smechr0302406 Solanum melongena
Solyc05g024260.3 Solanum lycopersicum
GLYMA_04G198500 Glycine max
HannXRQ_Chr09g0273741 Helianthus annuus
LOC111893962 Lactuca sativa
GSCOC_T00012727001 Coffea canephora
PGSC0003DMG400031738 Solanum tuberosum
C5167_006928 Papaver somniferum
0.50 0.40 0.30 0.20 0.10 0.00 Relative Times
Asparagaceae
Asteraceae
Brassicaceae
Bromeliaceae
Cannabaceae
Cucurbitaceae
Euphorbiaceae
Fabaceae
Linaceae
Malvaceae
Musaceae
Papaveraceae
Poaceae
Rosaceae
Rubiaceae
Rutaceae
Solanaceae
Vitaceae
Amaranthaceae
Arecaceae

**Supplementary Figure 6. Phylogenetic analysis for AtSWEET11 and AtSWEET12 orthologs in different plant species.**

The evolutionary analysis of **A,** *AtSWEET11* and **B,** *AtSWEET12* orthologs in thirty-nine different plant species from twenty different families was performed. The time-tree was generated using the RelTime method (1). Divergence times for all branching points in the topology were calculated using the Maximum Likelihood method and General Time Reversible model (2). The tree is drawn to scale with branch lengths measured in the relative number of substitutions per site. This analysis involved 25 nucleotide sequences for A, and 33 sequences for B. Evolutionary analysis were conducted in MEGA X (3). The name of the families for each species were indicated by colored circle. For more details see Supplementary File 1. For methodology details, please refer to the following literature:

1. Tamura K., Battistuzzi FU, Billing-Ross P, Murillo O, Filipski A, and Kumar S. (**2012**). Estimating Divergence Times in Large Molecular Phylogenies. *Proceedings of the National Academy of Sciences* **109**:19333-19338.
2. Nei M. and Kumar S. (**2000**). *Molecular Evolution and Phylogenetics*. Oxford University Press, New York.
3. Kumar S., Stecher G., Li M., Knyaz C., and Tamura K. (**2018**). MEGA X: Molecular Evolutionary Genetics Analysis across computing platforms. *Molecular Biology and Evolution* **35**:1547-1549.

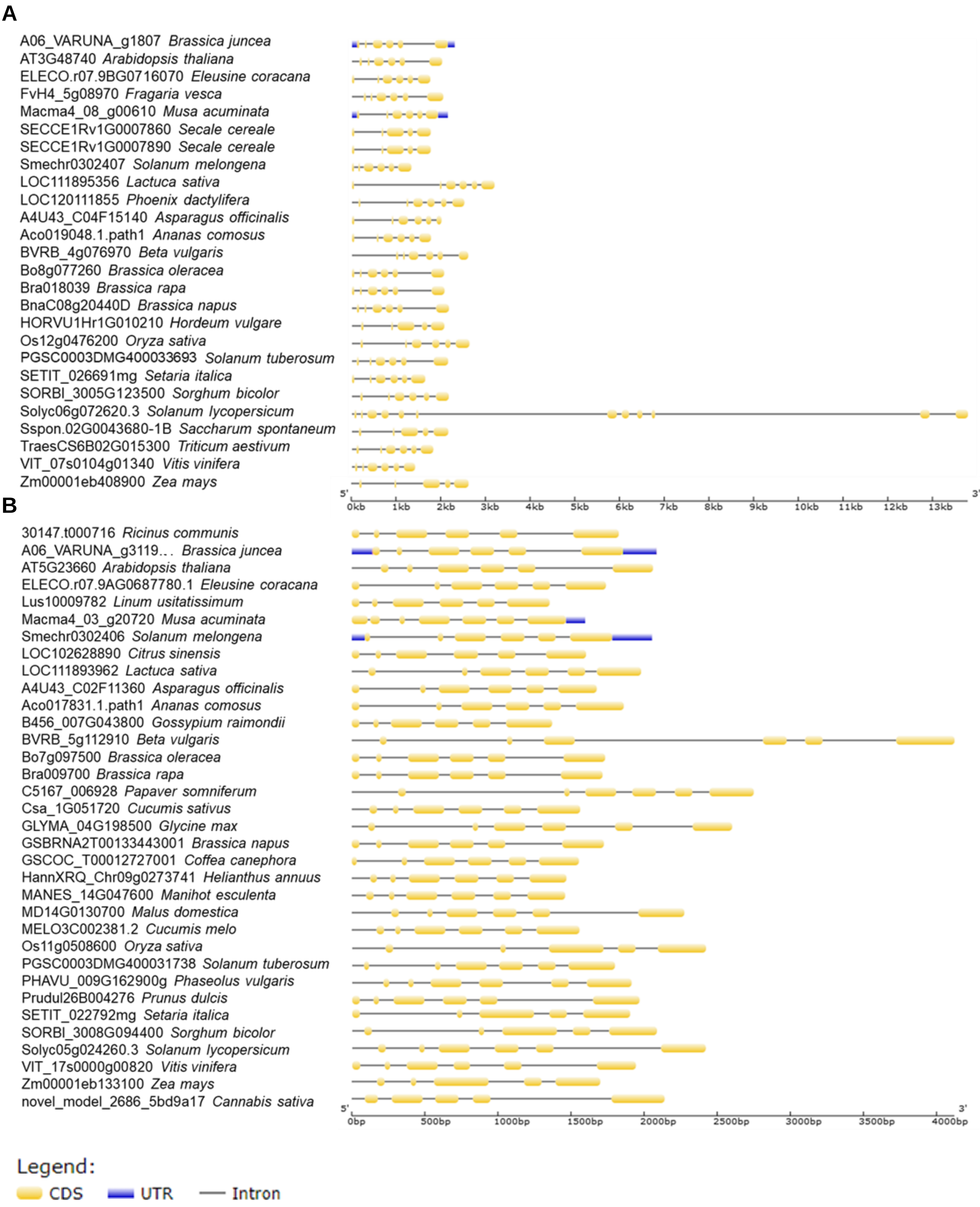
A
A06_VARUNA_g1807 Brassica juncea
AT3G48740 Arabidopsis thaliana
ELECO.r07.9BG0716070 Eleusine coracana
FvH4_5g08970 Fragaria vesca
Macma4_08_g00610 Musa acuminata
SECCE1Rv1G0007860 Secale cereale
SECCE1Rv1G0007890 Secale cereale
Smechr0302407 Solanum melongena
LOC111895356 Lactuca sativa
LOC120111855 Phoenix dactylifera
A4U43_C04F15140 Asparagus officinalis
Aco019048.1.path1 Ananas comosus
BVRB_4g076970 Beta vulgaris
Bo8g077260 Brassica oleracea
Bra018039 Brassica rapa
BnaC08g20440D Brassica napus
HORVU1Hr1G010210 Hordeum vulgare
Os12g0476200 Oryza sativa
PGSC0003DMG400033693 Solanum tuberosum
SETIT_026691mg Setaria italica
SORBI_3005G123500 Sorghum bicolor
Solyc06g072620.3 Solanum lycopersicum
Sspon.02G0043680-1B Saccharum spontaneum
TraesCS6B02G015300 Triticum aestivum
VIT_07s0104g01340 Vitis vinifera
Zm00001eb408900 Zea mays
5'
3'
0kb 1kb 2kb 3kb 4kb 5kb 6kb 7kb 8kb 9kb 10kb 11kb 12kb 13kb
B
30147.t000716 Ricinus communis
A06_VARUNA_g3119... Brassica juncea
AT5G23660 Arabidopsis thaliana
ELECO.r07.9AG0687780.1 Eleusine coracana
Lus10009782 Linum usitatissimum
Macma4_03_g20720 Musa acuminata
Smechr0302406 Solanum melongena
LOC102628890 Citrus sinensis
LOC111893962 Lactuca sativa
A4U43_C02F11360 Asparagus officinalis
Aco017831.1.path1 Ananas comosus
B456_007G043800 Gossypium raimondii
BVRB_5g112910 Beta vulgaris
Bo7g097500 Brassica oleracea
Bra009700 Brassica rapa
C5167_006928 Papaver somniferum
Csa_1G051720 Cucumis sativus
GLYMA_04G198500 Glycine max
GSBRNA2T00133443001 Brassica napus
GSCOC_T00012727001 Coffea canephora
HannXRQ_Chr09g0273741 Helianthus annuus
MANES_14G047600 Manihot esculenta
MD14G0130700 Malus domestica
MELO3C002381.2 Cucumis melo
Os11g0508600 Oryza sativa
PGSC0003DMG400031738 Solanum tuberosum
PHAVU_009G162900g Phaseolus vulgaris
Prudul26B004276 Prunus dulcis
SETIT_022792mg Setaria italica
SORBI_3008G094400 Sorghum bicolor
Solyc05g024260.3 Solanum lycopersicum
VIT_17s0000g00820 Vitis vinifera
Zm00001eb133100 Zea mays
novel_model_2686_5bd9a17 Cannabis sativa
5'
3'
0bp 500bp 1000bp 1500bp 2000bp 2500bp 3000bp 3500bp 4000bp
Legend:
CDS
UTR
Intron

**Supplementary Figure 7. Exon-intron distribution for *AtSWEET11* and *AtSWEET12* orthologs in different plant species.**

The gene structure analysis including exon-intron arrangement were conducted for the orthologs of **A,** *AtSWEET11* gene and **B,** AtSWEET12 gene from thirty-nine different plant species using Gene Structure Display Server 2.0 (http://gsds.gao-lab.org/index.php). The analysis involved 25 gene sequences for A, and 33 gene sequences for B. Exons are represented by yellow color; Introns are represented by black line; Untranslated-regions (UTRs) are represented by blue color. For more details see Supplementary File 1.

**A**

| | |
|---|---|
| *Brassica oleracea* _Bo8g077260 | 3 |
| *Arabidopsis thaliana* _AT3G48740 | 117 |
| *Oryza sativa* _Os12g0476200 | 61 |
| *Solanum lycopersicum* _Solyc06g072620.3 | 3 |
| *Solanum tuberosum* _PGSC0003DMT400083992 | 502 |
| *Setaria italica* _SETIT_026691mg | 0 |
| *Sorghum bicolor* _SORBI_3005G123500 | 3 |

| | |
|---|---|
| (white) | 0 |
| (light pink) | 10-50 |
| (pink) | 50-100 |
| (rose) | 101-200 |
| (dark red) | 201-1000 |
| (red) | >1000 |

**B**

| | |
|---|---|
| *Brassica oleracea* _Bo7g097500 | 1 |
| *Arabidopsis thaliana* _AT5G23660 | 25 |
| *Oryza sativa* _Os11g0508600 | 1 |
| *Solanum lycopersicum* _Solyc05g024260.3 | 37 |
| *Solanum tuberosum* _PGSC0003DMT400081206 | 0 |
| *Setaria italica* _SETIT_022792mg | 2262 |
| *Sorghum bicolor* _SORBI_3008G094400 | 179 |

**Supplementary Figure 8. Transcript expression of *AtSWEET11* and *AtSWEET12* orthologs in the leaf tissues from different plant species.**

**A and B,** The transcript expression pattern of **A,** *AtSWEET11* and **B,** *AtSWEET12* orthologs from *Brassica oleracea*, *Arabidopsis thaliana*, *Oryza sativa*, *Solanum lycopersicum*, *S. tuberosum*, *Triticum aestivum*, *Setaria italica* and *Sorghum bicolor* were obtained from publically available Expression Atlas (https://www.ebi.ac.uk/gxa/home). The transcript values were expressed as transcripts per millions (TPM) and were represented in the form of heat map. Color bar ranging from dark red to white indicate the levels of transcript expression from high to low or undetected respectively.

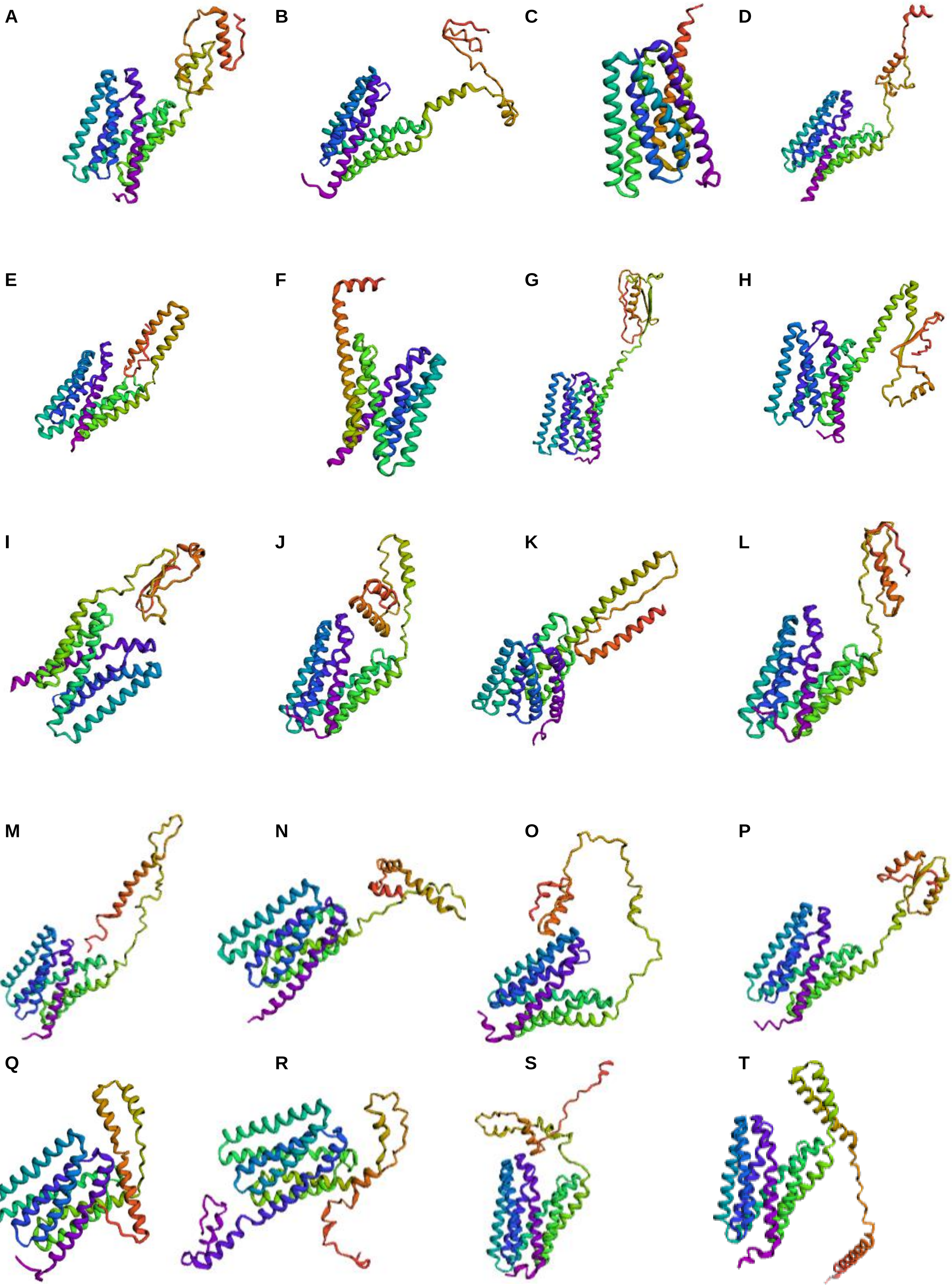
A
B
C
D
E
F
G
H
I
J
K
L
M
N
O
P
Q
R
S
T

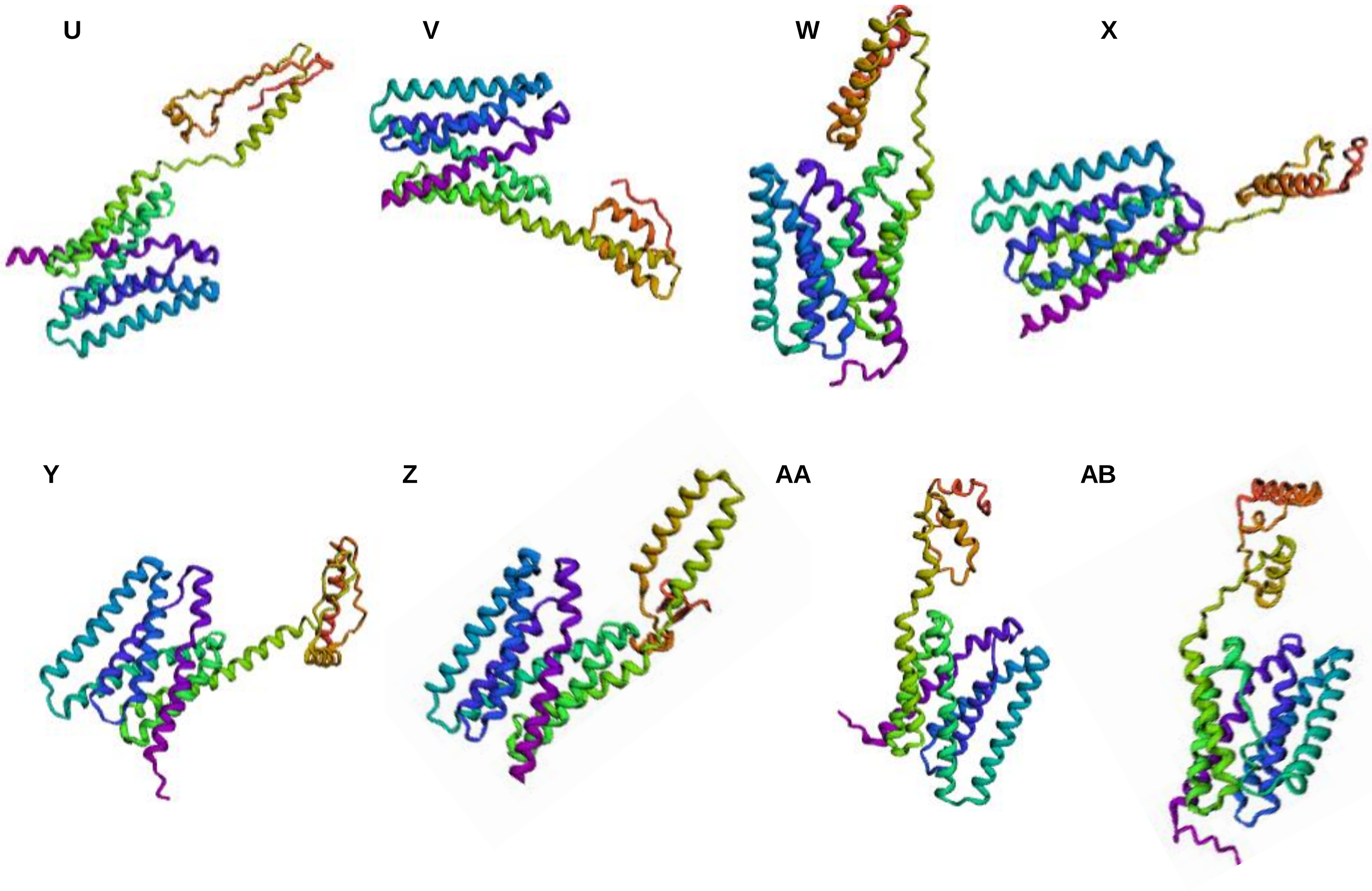

**Supplementary Figure 9. Tertiary structure prediction of AtSWEET11 and AtSWEE12 orthologs from different plants.** The tertiary structures of the proteins orthologus to SWEET11 and SWEET12 were predicted using Robetta (https://robetta.bakerlab.org/). The structures depicted in the figure represents a family A,B are from Brassicaceae C,D Asparagaceae E Asteraceae F,G Amaranthaceae H Cannabaceae I,J Bromeliaceae K Cucurbitaceae L Euphorbiaceae M Fabaceae N Linaceae

O Malvaceae P Papaveraceae Q,R Musaceae S,T Poaceae U Rosaceae V Rubiaceae W Rutaceae X Arecaceae Y,Z Solanaceae and AA, AB Vitaceae

This tertiary structure prediction analysis revealed that the majority of SWEET protein C-terminal regions are intrinsically disordered and the C-terminal region did not exhibits any ordered structure.

**A**

**SWEET11**

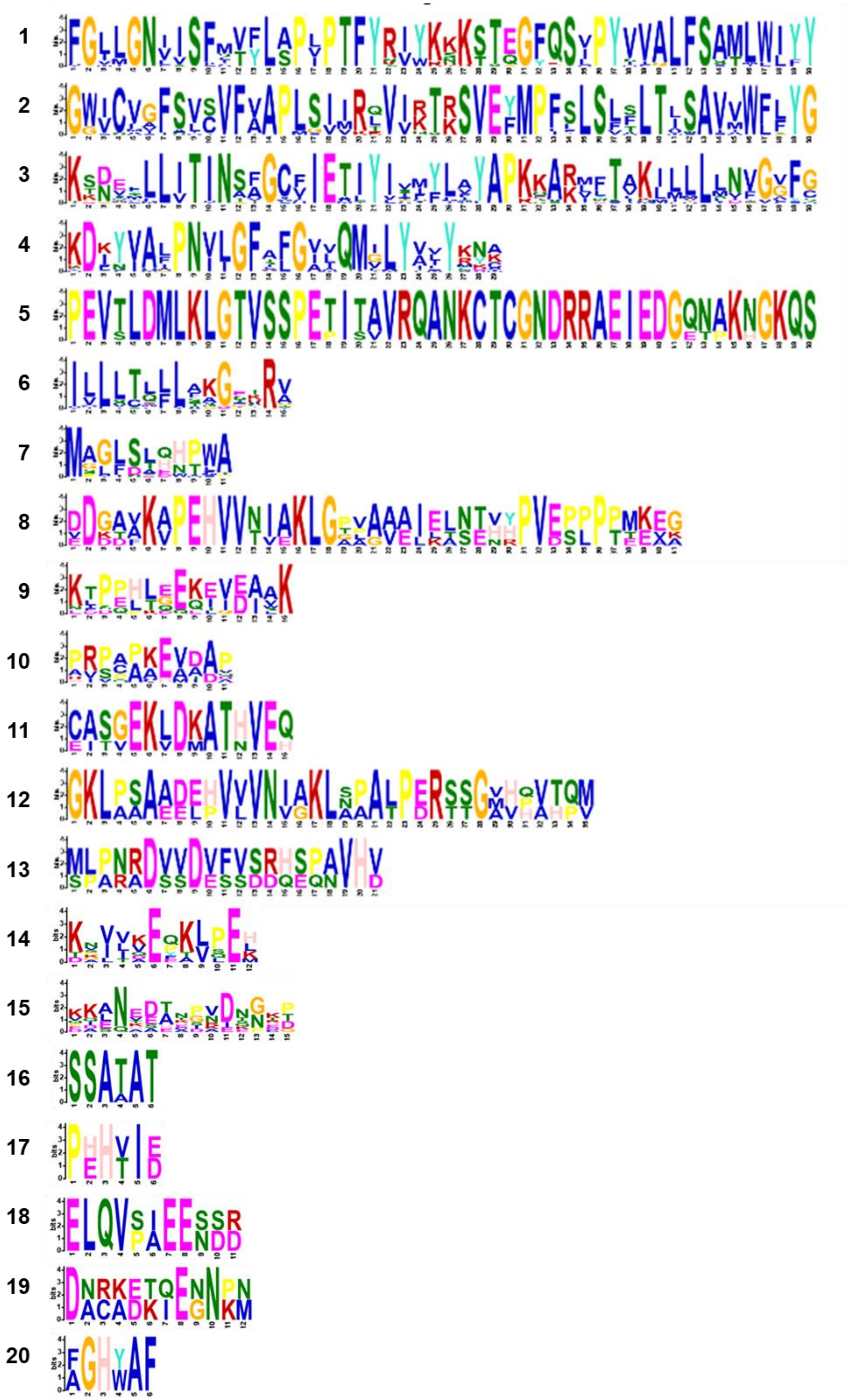

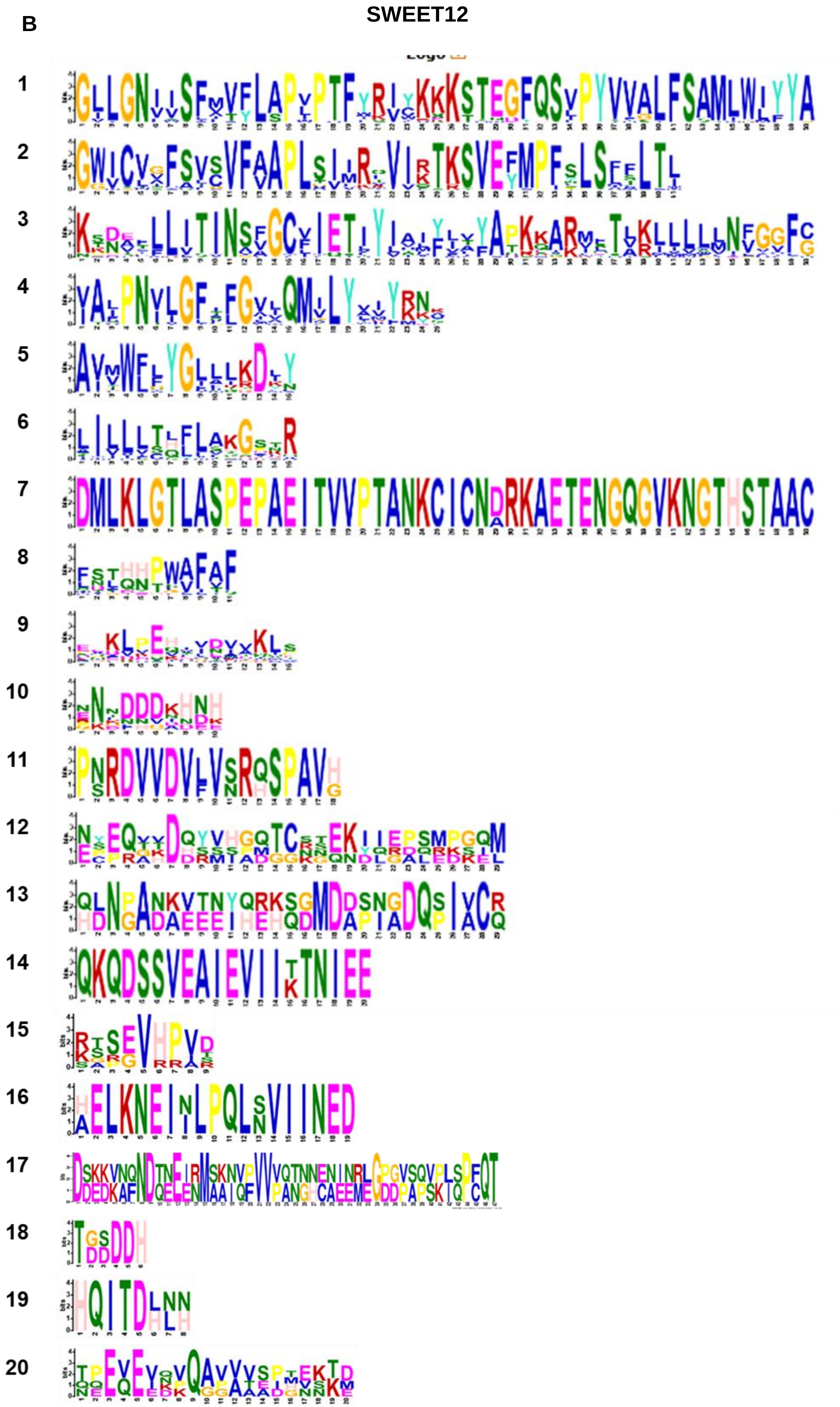
B
SWEET12

**Supplementary Figure 10. Details of logo of each protein motifs for AtSWEET11 and AtSWEET12 orthologs in different plant species.**

**A and B**, The logo of each motif and associated amino acids were identified for **A,** AtSWEET11 and **B,** AtSWEET12 protein orthologs from different plant species using MEME_suite (https://meme-suite.org/meme/). The amino acid frequencies and level of their conservation were indicated by the heights of letters. The X-axis indicates the length of motif and Y-axis indicates the sequence conservation per site of each letter (i.e., bit score). The numbering of the motifs was done according to its significance, the highly significant motifs i.e. the first four motifs were conserved in most of the sequences used in present study. These motifs corresponds to the transmembrane regions of the proteins, conferring that these orthologous proteins could be performing similar functions.

A

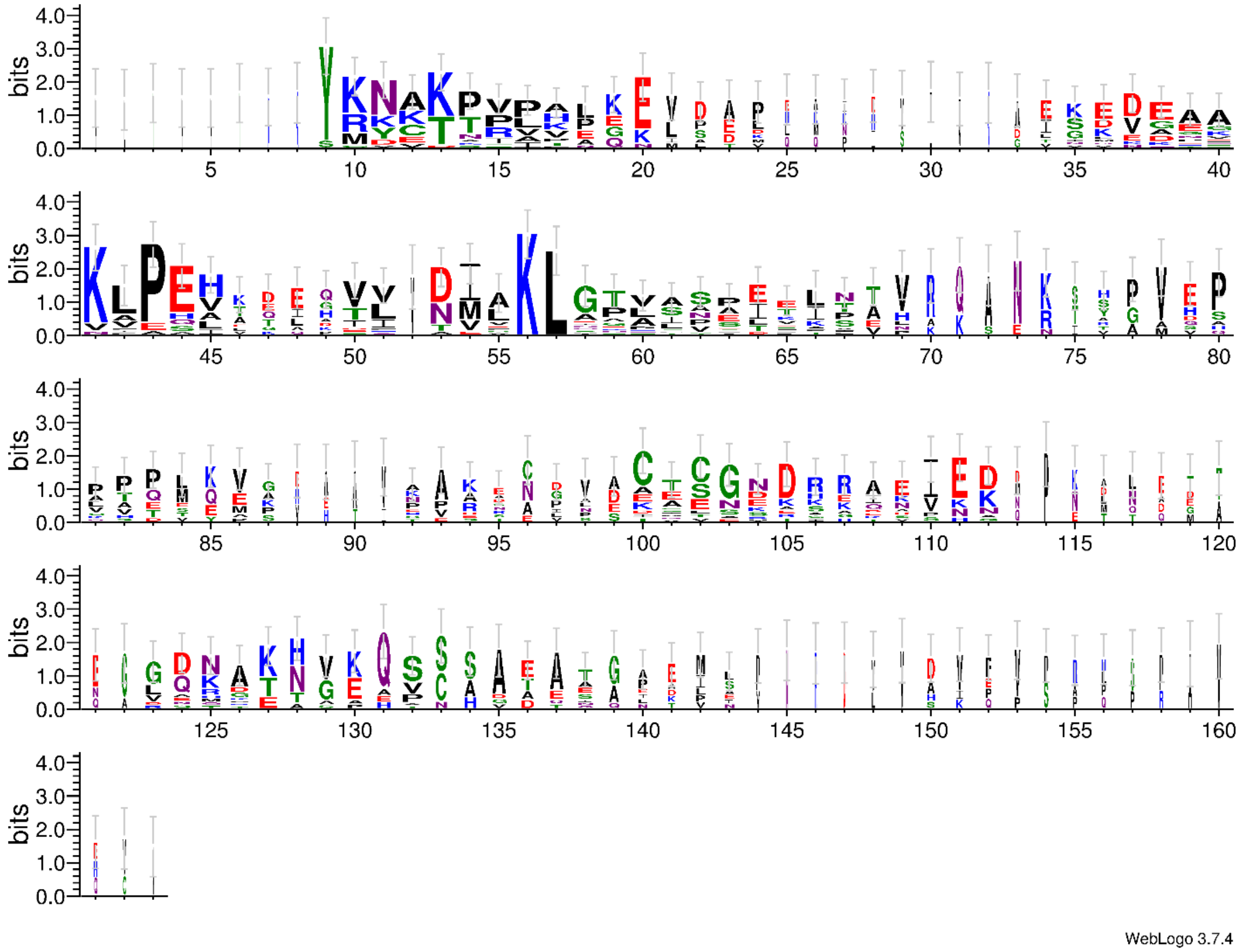

B

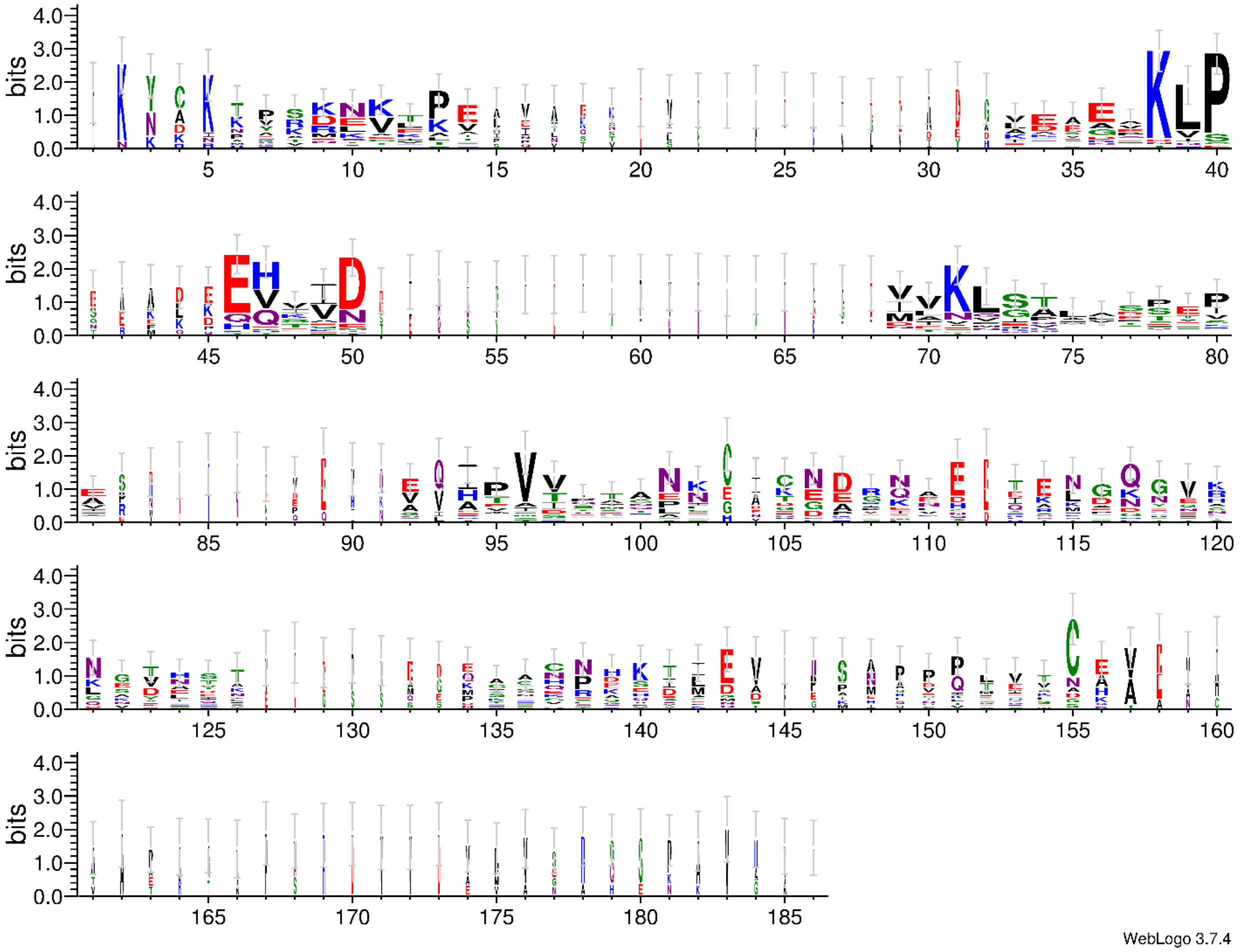

**Supplementary Figure 11. The C-terminal analysis of AtSWEET11 and AtSWEET12 protein orthologs from different plant species.**

The amino acids sequences from C-terminal of **A,** AtSWEET11 and **B,** AtSWEET12 protein orthologs from thirty-nine different plant species were extracted and aligned using the Clustal Omega (https://www.ebi.ac.uk/Tools/msa/clustalo/). The logo of associated amino acids were generated using WebLogo3 (http://weblogo.threeplusone.com/).The amino acid frequencies and level of their conservation were indicated by the heights of letters. The Y-axis indicates the sequence conservation per site of each letter (i.e., bit score). The complete sequence for AtSWEET11 and AtSWEET12 proteins were obtained from ensemble plants/Phytozome. These sequences were used as input for TMHMM to determine the co-ordinates of the C-terminal sequences. Using an in-house perl script, the resulting co-ordinates were then used to extract C-terminal sequences.

**Supplementary Table 1: Sucrose interacting residues in AtSWEET11 and AtSWEET12 with equivalent dCMP binding residues in AtSWEET13 crystal structure.**

!

| AtSWEET11 | AtSWEET12 | AtSWEET13 (ligand binding pocket: PDB: 5XPD) (1) |
|---|---|---|
| S22 | S22 | S20 |
| S56 | S56 | S54* |
| W60 | W60 | W58 |
| N77 | N77 | N76 |
| N197 | N197 | N196 |
| S143 | S143 | S142 |
| S177 | S177 | S176* |
| W181 | W181 | W180 |
| | | V23* |
| V146 | V146 | V145* |
| F26 | F26 | |
| A53 | A53 | |
| A57 | A57 | |
| L73 | L73 | |
| I76 | I76 | |
| F147 | F147 | |
| L174 | | |
| A178 | | |
| P196 | P196 | |
| G200 | | |

* asterisk indicates mutated residues in AtSWEET13-dCMP crystal structure.

!

**Supplementary Table 2: The details of conserved residues among AtSWEETs and OsSWEET2b.**

| Conserved residues among AtSWEET11 | AtSWEET12 | AtSWEET13 | OsSWEET2b |
|---|---|---|---|
| G15 | G15 | G13 | G13 |
| G18 | G18 | G16 | G16 |
| N19 | N19 | N17 | N17 |
| P29 | P29 | P27 | P27 |
| T32 | T32 | T30 | T30 |
| P49 | P49 | P47 | P47 |
| Y50 | Y50 | Y48 | Y48 |
| Y63 | Y63 | Y61 | Y61 |
| G136 | G136 | G135 | G136 |
| P150 | P150 | P149 | P150 |
| V157 | V157 | V156 | V157 |
| T160 | T160 | T159 | S160 |
| S162 | S162 | S161 | S162 |
| M166 | M166 | M165 | M166 |
| P167 | P167 | P166 | P167 |
| F168 | F168 | F167 | F168 |
| L170 | L170 | L169 | L170 |

| | | | |
|---|---|---|---|
| S171 | S171 | S170 | S171 |
| Y184 | Y184 | Y183 | Y184 |
| D190 | D190 | D189 | D190 |
| N197 | N197 | N196 | N197 |
| Q207 | Q207 | Q206 | Q207 |

The twenty-two conserved residues among AtSWEETs and OsSWEET2b.

T159 (AtSWEET13) is conserved among all AtSWEEts, while S160 is present in OsSWEET2b. N197 is also a part of substrate binding pocket as depicted via AtSWEET13 crystal structure.
